# Plastid phylogenomics and cytonuclear discordance in Rubioideae, Rubiaceae

**DOI:** 10.1101/2024.01.17.576054

**Authors:** Olle Thureborn, Niklas Wikström, Sylvain G. Razafimandimbison, Catarina Rydin

## Abstract

In this study of evolutionary relationships in the subfamily Rubioideae (Rubiaceae), we take advantage of the off-target proportion of reads generated via previous target capture sequencing projects based on nuclear genomic data to build a plastome phylogeny and investigate cytonuclear discordance. The assembly of off-target reads resulted in a comprehensive plastome dataset and robust inference of phylogenetic relationships, where most intratribal and intertribal relationships are resolved with strong support. While the phylogenetic results were mostly in agreement with previous studies based on plastome data, novel relationships in the plastid perspective were also detected. For example, our analyses of plastome data provide strong support for the SCOUT clade and its sister relationship to the remaining members of the subfamily, which differs from previous results based on plastid data but agrees with recent results based on nuclear genomic data. However, several instances of highly supported cytonuclear discordance were identified across the Rubioideae phylogeny. Coalescent simulation analysis indicates that while ILS could, by itself, explain the majority of the discordant relationships, plastome introgression may be the better explanation in some cases. Our study further indicates that plastomes across the Rubioideae are, with few exceptions, highly conserved and mainly conform to the structure, gene content, and gene order present in the majority of the flowering plants.

## Introduction

The nuclear genome, the plastid genome (plastome) and the mitochondrial genome (mitogenome) are the three genomes in plant cells. The high number of copies in the cells, generally clonal and uniparental inheritance, and highly conserved structure, arrangement and gene content have made the plastome the most utilized genome for molecular phylogenetic analyses at different taxonomic ranks among green plants [1–3]. A drawback of relying solely on information from the plastome or mitogenome for phylogenetic reconstruction is that, due to their mode of inheritance, results based on one of the organellar genomes are generally considered to represent only a single gene tree within the species tree [3, 4]. Hence, they may provide misleading results on species relationships in case of for example incomplete lineage sorting (ILS; i.e., temporal mismatch between gene divergence and species divergence resulting in the retention of ancestral gene polymorphism in the genome) or hybridization [3–6]. Lack of sufficient phylogenetic information in commonly used plastid markers has also led to unsatisfactory resolution, at least in studies of closely related species. The spacers of the nuclear ribosomal DNA have often proven informative and have been frequently utilized in phylogenetic studies of plants, often in combination with plastid markers in order to increase the amount of informative characters. However, since these spacers (e.g., the much-used internal transcribed spacer, nrITS) are relatively short, at least in angiosperms [7, 8], they can nevertheless only contribute a limited amount of information.

With recent advances in molecular and computational methods, the generation and analysis of genome-scale datasets have become feasible. High-throughput sequencing of nuclear genomes allow analysis of a large collection of unlinked genes, which should increase the chance of accurately reconstruct the species tree [5, 9–11]. Sequencing of complete or near-complete plastomes has also become routine and the increased amount of data has resulted in better supported plastid phylogenies [12, 13]. Contrasting nuclear and organellar trees (i.e., phylogenies produced from nuclear DNA data, and plastid/mitochondrial DNA data, respectively) may reveal conflicting phylogenetic patterns (cytonuclear discordance) [6, 14–16]. While methodological artifacts due to for example sparse taxon sampling can result in such conflicts [17], cytonuclear discordance is often attributed to real biological processes such as ILS or hybridization and introgression (backcrossing of a hybrid with a parent species) leading to chloroplast capture (fixation of the chloroplast genome of one lineage in another lineage) [1, 14, 18–21]. Although cytonuclear discordance is often detected, the possibility to distinguish between ILS and hybridization processes/chloroplast capture has been limited as currently available methods are dependent on large samples of nuclear markers [20, 22].

While still not commonplace, the production of phylogenomic nuclear datasets are expected to rapidly increase due to the availability of universal target capture kits [23], such as the Angiosperms353 kit [24]. Target capture (or target enrichment) selects for specific genomic regions of interest using baits that are designed to fit the study group based on information in reference sequences (e.g., transcriptomes or whole genomes) from related species [24–26]. In this way more samples can be sequenced in the same sequencing run and financial as well as computational costs are substantially decreased compared to transcriptome and whole genome sequencing [24–26]. While genome skimming, or shallow whole genome sequencing [27] is the main approach for generating plastome data [28–30], the portion of off target reads when using a target capture approach can also be sufficient to recover at least large portions of the plastome [31–33].

The subfamily Rubioideae, currently with 29 tribes and about 8000 species, is the largest of the subfamilies of the coffee family (Rubiaceae), comprising about 58% of the species [34–36]. Previous phylogenomic estimates of this group include the tribal-level phylogenies of Rubiaceae [35, 37] and Gentianales [38]. While Rydin et al. [37] and Wikström et al. [35] used genome skimming to obtain mitochondrial and nuclear ribosomal cistron + plastome data, Antonelli et al. [38] used target capture sequencing to obtain 353 nuclear genes targeted with the Angiosperms353 kit and partial plastomes. Although corroborating and clarifying several parts of the backbone of the Rubioideae phylogeny, uncertainty regarding several relationships remained, either as a result of low support, discordance, or sparse taxon sampling.

Recently, the Rubioideae were the focus of a taxon-rich phylogenomic analysis based on target capture nuclear data recovered by using the Angiosperms353 kit [36]. That study closed important taxon gaps and expanded the number of representatives per tribe and characters considerably and resulted in a comprehensive nuclear phylogenetic hypothesis of the Rubioideae with almost all relationships highly supported in both coalescent-based and concatenation-based trees. The pattern of gene tree conflict at short internodes suggested that rapid speciation events have been relatively common in the group and is a likely reason why some parts of the phylogeny had remained difficult to resolve. While the obtained nuclear results often corroborated those of previous studies, some highly supported conflicts with results in previous studies mostly based on plastid data were also noted. Although those cytonuclear conflicts may well be due to biological causes, this should preferably be tested using identical sets of taxa for plastome and nuclear data.

Here we utilize off-target plastid reads to assemble a large dataset of the Rubiaceae subfamily Rubioideae. Our taxon sampling is greatly increased compared to previous studies of the subfamily based on plastome data, and is identical with the taxon sampling used in Thureborn et al. [36]. Using phylogenomic analyses our main goals were to (1) clarify the plastome phylogeny of the Rubioideae, and (2) examine whether ILS or hybridization/introgression best explains any detected cases of cytonuclear discordance within the group. A secondary goal was to gain a better understanding of plastome evolution within the Rubioideae.

## Materials and methods

### Taxon sampling

The aim was to employ a taxon sampling identical to that of the nuclear target capture data study by Thureborn et al. [36]. The same specimens used in that study were thus used here. The dataset included a total of 124 species, of which 101 were Rubioideae species and 23 were outgroup species (S1 Table). Within the Rubioideae 27 of the 29 commonly recognized tribes were represented. The vast majority of tribes were represented by more than one species. The two unsampled tribes are Foonchewieae and the recently described Aitchisonieae, both monospecific. Outgroup taxa included 20 species from other major lineages of Rubiaceae as well as three non-Rubiaceae species from the gentianalean families Gentianaceae, Loganiaceae and Apocynaceae. For four specimens (*Colletoecema dewevrei*, *Mitchella repens*, *Prismatomeris fragrans* and *Schizocolea linderi*) plastome sequences had already been produced [35] and were downloaded from GenBank. For the other specimens the corresponding target capture sequencing reads that were used in Thureborn et al. [36] were reused. For one specimen (*Leptostigma pilosum*), the target capture reads were complemented with genome skimming reads available from another sequencing project (Thureborn et al., unpublished data).

The production of genome skimming reads included extraction of genomic DNA using a CTAB protocol [39] followed by library preparation and high-throughput sequencing carried out by Science for Life Laboratory (Solna, Sweden). Library inserts (aiming at 350 bp) were prepared using the ThruPLEX DNA-seq library preparation kit (Rubicon Genomics, Ann Arbor, USA). The library was sequenced together with 92 other libraries on the Illumina HiSeq2500 platform (Illumina, San Diego, USA) with 2 x 126 bp paired-end reads. Target capture sequencing reads for 93 of the 124 species were generated by Thureborn et al. [36] and the other 31 species were available through the Plant and Fungal Tree of Life (PAFTOL) Research Program [23]. In brief, target capture reads were generated using a target capture approach, utilizing the universal Angiosperms353 probe set [24], coupled with Illumina sequencing. For more details on the generation of the targeted sequencing reads see Thureborn et al. [36] and Baker et al. [23]. A list of all included taxa including voucher and GenBank/ENA information is found in S1 Table.

### Sequence data preprocessing and plastome assembly

The processed reads used in Thureborn et al. [36] were reused here. Preprocessing of reads were conducted using tools of the BBtools package [40] and included adapter removal, quality trimming (Q<20) and removal of short reads (<36bp) using BBDuk v37.64, and deduplication using dedupe v37.64 or alternatively clumpify v38.90. BBDuk and dedupe were used as implemented in Geneious v11.1.5 [41]. Genome skim reads were processed the same way. The total number of cleaned reads per sample ranged between 424 886 and 61 673 872 (mean = 12 372 381) (S1 Table).

Plastomes were assembled using either a reference-guided approach, a *de novo* approach, or a combined reference-guided + *de novo* approach. An initial mapping of each sample to a closely related reference plastome sequence indicated good coverage and contiguity for some samples whereas other samples had relatively low coverage and contiguity. Assembly was initially conducted using the NOVOWrap v1.20 platform [42], which internally used the seed-extend based *de novo* assembler NOVOPlasty v3.7.2 [43], with *Coffea arabica* (GenBank accession NC_008535) set as reference. Nineteen circularized plastomes were obtained using this method.

Another 70 samples that were not successfully assembled using NOVOWrap were instead assembled using a combination of reference mapping and *de novo* assembly. Reads from each of those samples were mapped to a reference plastome sequence of a closely related species using the Geneious read mapper with medium-low sensitivity for up to five iterations. The mapped reads were then *de novo* assembled using Spades v3.10.0 [44] in careful mode.

Contigs identified as originating from the plastome were extended by iterative mapping using the Geneious read mapper. Contigs were manually inspected and edited if necessary (mainly by trimming questionable sequence from their ends). The Geneious assembler was used to merge overlapping contigs. When this procedure did not produce one single contig, the contigs were instead ordered according to the reference by mapping or using Mauve v2.3.1 [45] with the Mauve Contig Mover algorithm [46] with *Coffea arabica* (GenBank accession NC_008535) set as reference genome and adding a spacer of 100 Ns between each contig. Draft plastomes were subjected to further iterative mapping using the Geneious mapper to identify and correct errors. Occasionally, local re-assemblies were conducted to verify problematic regions, using Spades and/or the Geneious assembler. To verify the assemblies (including the NOVOWrap/*de novo* assemblies), a final round of mapping was conducted with medium-low sensitivity for up to five iterations and restricting paired reads to map nearby, and contigs were manually inspected for errors and edited if necessary. Gaps in consensus sequences were coded as Ns.

Thirty-one species were subject to reference-guided assembly using the Geneious mapper with medium-low sensitivity for up to five iterations and restricting paired reads to map nearby. Consensus sequences were trimmed to the reference (which had one IR removed) and gaps were coded as Ns. Plastome sequences were annotated using GeSeq v2.03 [47] and Geneious (including manual adjustments). Boundaries of the inverted repeat (IR) region were identified searching for repeats using the Find repeats tool implemented in Geneious.

### Alignment and phylogeny

During assembly and initial alignment large inversions were detected in four of the assembled plastomes. The endpoints of the inversions were checked by individually aligning each of the four plastomes with *Coffea arabica* (GenBank accession NC_008535) set as reference using Mauve with the progressiveMauve algorithm [48]. The inversions were reverse complemented before whole plastome alignment with MAFFT v7.450 as implemented in Geneious, using the FFT-NS-x 1000 algorithm (gap opening penalty set to 1.53; offset value set to 0.0). Only one IR region was included in the alignments to avoid data duplication.

Using alignment trimming tools to remove ambiguously aligned columns or missing data prior to phylogenetic inference have had mixed success on phylogenetic accuracy, with untrimmed or softly trimmed alignments often performing just as well or better than more aggressively trimmed alignments [49–51]. Here, to test the robustness of our phylogenetic results we employed two different trimming strategies. Columns containing only missing data characters (“N” and “-“ characters) and autapomorphic insertions were deleted from the alignment by first converting “N”s to “-“ characters, followed by trimming columns where more than 99% of the samples had a gap (alignment version 1), using the Mask alignment tool in Geneious. Another alignment (alignment version 2) was created by trimming the alignment using trimAl v1.2rev57 [52] with the gappyout parameter (chosen by the heuristic automated1 method).

Additionally we applied RY-coding to the two alignment versions using BMGE v1.12 [53], thus resulting in a total of four alignment versions. This method recodes the four nucleotides into two state categories, purines (A,G = R) and pyrimidines (C,T = Y), and is used to limit the effect of substitution saturation and compositional bias, although it comes at the expense of decreased information content in the dataset [54–58].

IQ-TREE 2 v2.1.2 [59] was used via the Cipres Science Gateway [60] to infer a maximum likelihood tree for each of the four alignment versions (alignments 1 and 2, and alignments 1 and 2 employing RY-coding). We ran unpartitioned analyses with automatic model selection [61]; the selected model was the TVM + F + I + G4 model for the non-RY coded alignments, and TVMe+I+G4 for the RY-coded alignments. Branch support was assessed with 1000 ultrafast bootstrap replicates applying the hill-climbing nearest-neighbor interchange search to guard against overestimation of support [62].

### Coalescent simulation analyses

To evaluate if discordance between plastid and nuclear phylogenies could be attributed to ILS alone we followed an approach used in several previous studies [16, 20, 21, 31, 63]. One thousand plastome trees were simulated under a neutral coalescent model using the contained_coalescent_tree function in DendroPy v4.5.2 [64]. As guide tree we used the ASTRAL species tree based on supercontigs (coding + noncoding data) of 353 nuclear genes in Thureborn et al. [36]. Generally, under the assumptions of haploidy and uniparental inheritance of organellar genomes, the effective population size of organelle genomes in hermaphroditic and dioecious taxa are expected to be half and one fourth of the (diploid) nuclear genome, respectively [1]. Or put differently, the coalescence rate of the plastome is expected to be two times (hermaphroditic taxa) or four times (dioecious taxa) higher relative to that of the nuclear genome. Therefore, to account for organellar inheritance, the branch lengths (in coalescent units) of the species tree (the guide tree) were rescaled by a factor of two and four using Gotree v0.4.1 [65] before simulations. IQTREE 2 was used to summarize the fraction of the simulated trees concordant with each branch of the observed plastome tree. If cytonuclear conflicts are due to ILS alone, a sizeable fraction of the observed plastome relationships that are in conflict with those of the nuclear guide tree should be present in the simulated plastome trees; conversely, if the observed plastome relationships are at or close to zero in frequency in the simulated trees, a hybridization scenario followed by chloroplast capture may be invoked [20, 21, 31]. Here we considered the hypothesis of ILS unlikely if the frequency of the observed plastome relationship was less than or equal to 1%.

## Results

### Dataset characteristics

The final alignment version 1 was 182 216 bp in length, and the final alignment version 2 was 128 382 bp in length (Table 1). The average degapped length of the individual sequences in alignment version 1 was 122 296 bp (range 13 342–130 482 bp), and 118 991 bp (range 13 419–126 621) in alignment version 2 (Table 1). Across both alignments, five species had sequences shorter than 75 000 bp: *Cyanoneuron pedunculatum* (13 442–13 319 bp), *Kajewskiella trichantha* (38 236–37 675 bp), *Jackiopsis ornata* (54 710–53 564 bp), *Chaetostachydium barbatum* (70 061–68 957 bp), and *Psychotria pandurata* (74 365–72 994 bp). These samples also had a relatively low amount of trimmed and deduplicated reads (S1 Table), which is likely a major reason for the poor sequence recovery. Across both datasets, 116 species had sequences longer than 100 000 bp.

**Table 1.** Characteristics of the two alignments used for phylogenetic inference.

| Align<br>ment | Trimming | Alignment<br>length | # of PIS | Mean degapped sequence<br>length (min-max) | Evolutionary<br>model (RY coded) |
| --- | --- | --- | --- | --- | --- |
| v. 1 | 99% gaps threshold /<br>autapomorphic<br>insertions | 182 216 | 58 390 | 122 296 (13 342–130 482) | TVM+F+I+G4<br>(TVMe+I+G4) |
| v. 2 | trimAl (gappyout) | 128 382 | 48 653 | 118 911 (13 419–126 621) | TVM+F+I+G4<br>(TVMe+I+G4) |

### Plastome phylogeny

The Rubioideae phylogenies inferred from all alignment versions were highly similar in both topology and support values, with strong bootstrap support (BS), 95% or more, for the vast majority of nodes (non-RY coding: Figs 1 and 2; and RY-coding: S1 and S2 Figs). Only a few nodes had lower bootstrap support than 100%. Compared to the corresponding non-RY coded tree, RY-coding only affected the poorly supported placement of the Colletoecemateae-Seychelleeae clade in the analysis of alignment 2 (S2 Fig). Support values remained high in most cases although some decreases in clade support led to fewer highly supported branches in the trees based on RY-coded data compared to the corresponding non-RY coded tree. For example, the SCOUT clade was supported by 97% bootstrap based on analysis of alignment 1 (Fig 1) and 99% based on alignment 2 (Fig 2), but support decreased to 69% (S1 Fig) and 94% (S2 Fig) in respective analysis of the RY-coded versions of these two alignments. Similarly, support for the Argostemmateae-Paederieae-Putorieae-Rubieae-Theligoneae clade decreased from BS=96% using non-RY coding of alignment 1 (Fig 1) to 93% when RY-coding was employed (S1 Fig). As the non-RY-coded and RY-coded results were highly similar and some decrease in support values is expected when applying RY-coding we focus on the results obtained with the non-RY-coded alignment versions (Figs 1 and 2).

**Fig 1.**
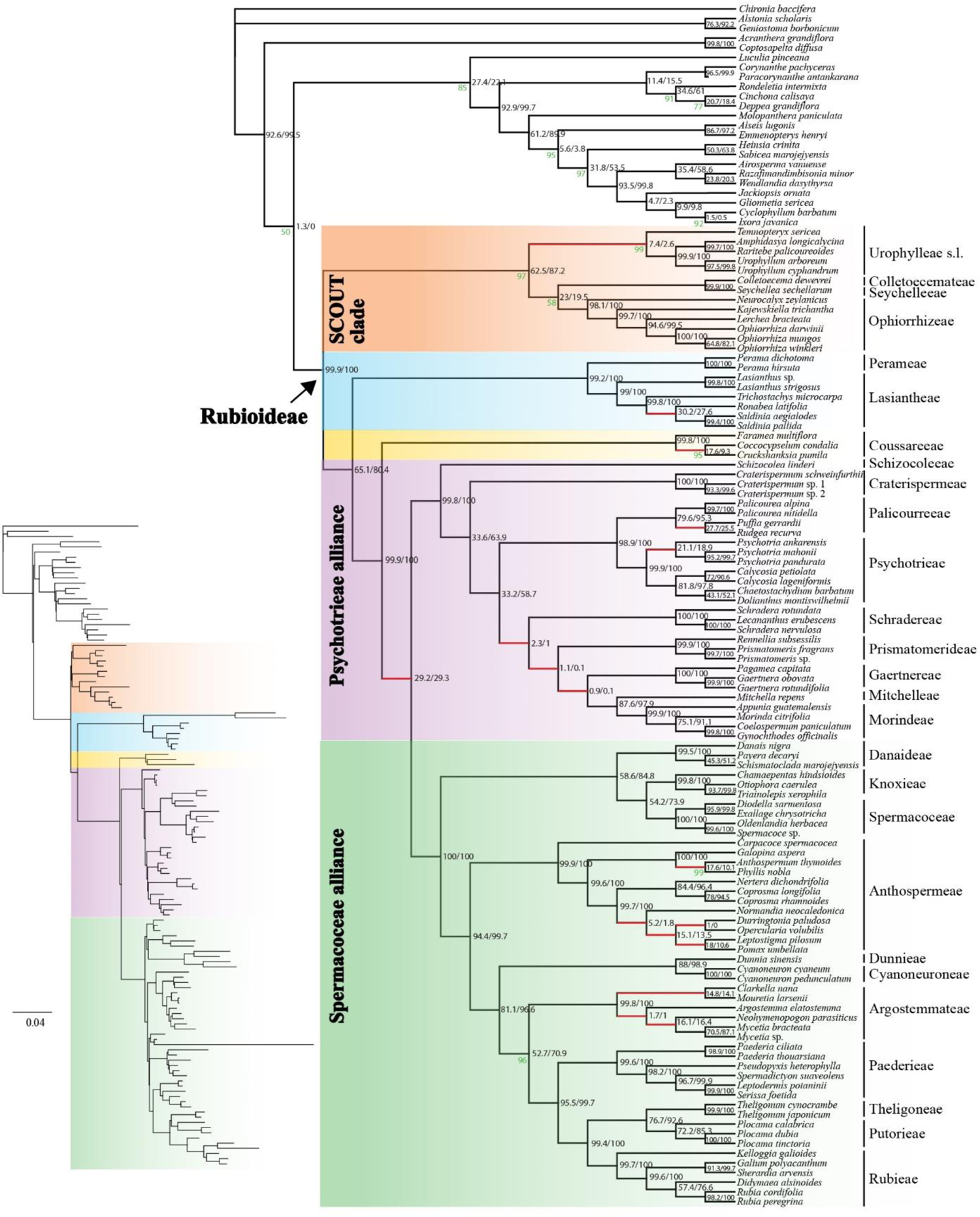
Plastome phylogeny inferred from the maximum likelihood analysis of the alignment version 1. The dataset was untrimmed except for removal of autapomorphies (columns where more than 99% of the samples had a gap). Support values are 100% unless otherwise indicated (in green). Values (in black) indicate the proportion of the 1000 simulated plastome trees (x2 scalar/x4 scalar) supporting the observed plastome relationship. Red branches indicate supported (≥95%) topological conflict with the nuclear coalescent-based and/or concatenation-based tree(s) in Thureborn et al. [36]. Inset branch lengths indicate the number of expected substitutions per site.

**Fig 2.**
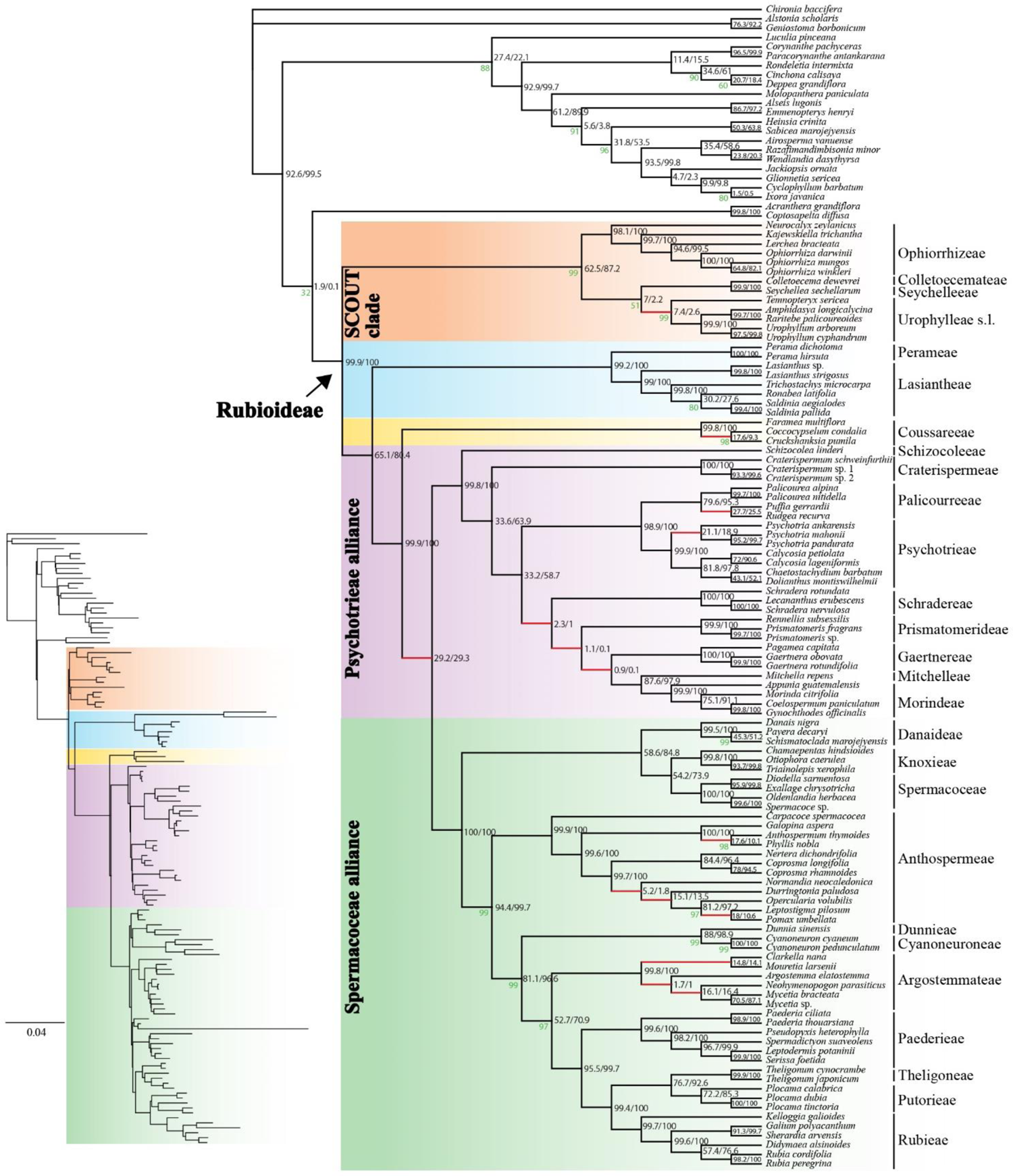
Plastome phylogeny inferred from the maximum likelihood analysis of the alignment version 2. The dataset was trimmed using trimAl. Support values are 100% unless otherwise indicated (in green). Values (in black) indicate the proportion of the 1000 simulated plastome trees (x2 scalar/x4 scalar) supporting the observed plastome relationship. Red branches indicate supported (≥95%) topological conflict with the nuclear coalescent-based and/or concatenation-based tree(s) in Thureborn et al. [36]. Inset branch lengths indicate the number of expected substitutions per site.

Concerning the phylogeny of the Rubioideae, only one node (alignment 1; Fig 1) and two nodes (alignment 2; Fig 2), respectively, were not strongly supported. The only strongly supported topological conflict between analyses of the two alignments was that *Durringtonia* and *Opercularia* formed a clade (BS = 100%) and were together sister clade to the *Leptostigma – Pomax* clade (BS = 100%) in the analysis of the alignment 1 (Fig 1), whereas *Durringtonia* was resolved as sister to an *Opercularia* + *Leptostigma – Pomax* clade (BS = 97%) in the analysis of alignment 2 (Fig 2). The second conflict was the poorly supported position of the Colletoecemateae *–* Seychelleeae clade, which was sister to Ophiorrhizeae (BS = 58%) based on analysis of the alignment 1 (Fig 1) but to Urophylleae s.l. (sensu Smedmark et al. [66], i.e., including *Temnopteryx*) (BS = 51%) in analysis of the alignment 2 (Fig 2). Apart from those differences the two trees were identical in topology. Results based on the alignment 1 had one more highly supported branch compared to those of the alignment 2, which was due to the differing level in support for the *Ronabea* + *Saldinia* clade (BS = 100% vs. 80%; Figs 1-2) within the tribe Lasiantheae. The subfamily Rubioideae, the Spermacoceae and Psychotrieae alliances, and all tribes were recovered as monophyletic (Figs 1 and 2).

The sister group to the remaining Rubioideae was composed of Colletoecemateae, Ophiorrhizeae, Seychelleeae, and Urophylleae s.l. This clade had the same constituents as the SCOUT clade in Thureborn et al. [36] but apart from the well-supported Seychelleeae *–* Colletoecemateae clade, intertribal relationships within the SCOUT clade are not well supported and differ between the alignment versions (see also above).

The SCOUT clade was sister to a clade formed by the Lasiantheae *–* Perameae clade, the tribe Coussareeae and a lineage that unites the Psychotrieae and Spermacoceae alliances (Figs 1 and 2).

Within the Psychotrieae alliance, Schizocoleeae and Craterispermeae were consecutive sisters to the remaining members of the alliance. The remaining members were divided into a clade uniting Psychotrieae + Palicoureeae and a clade formed by Schradereae, Prismatomerideae, Gaertnereae, Morindeae and Mitchelleae (Figs 1 and 2). Within the latter clade, Schradereae and Prismatomerideae are successive sisters to a Gaertnereae + Morindeae *–* Mitchelleae clade (Figs 1 and 2).

In the Spermacoceae alliance, a clade that unites Danaideae and Spermacoceae + Knoxieae were together sister to the remaining tribes, followed by Anthospermeae, a clade that unites Dunnieae + Cyanoneuroneae, Argostemmateae, Paederieae, a Theligoneae + Putorieae clade, and Rubieae (Figs 1-2).

### Cytonuclear conflicts

We identified some strongly supported conflicting relationships between our reconstructed plastome relationships and the recently published phylogenetic results based on nuclear data [36], i.e., the phylogenetic results based on supercontigs (coding and noncoding data) of 353 nuclear genes (figures 1 and 2 in Thureborn et al. [36]). While most intertribal relationships were concordant among plastid- and nuclear-derived phylogenies, the placements of Gaertnereae and Coussareeae differed between datasets from the different genomic compartments; Gaertnereae is here sister to Mitchelleae *+* Morindeae (Figs 1-2) but was sister to Palicoureeae *+* Psychotrieae based on nuclear data [36], and Coussareeae is here sister to the Psychotrieae and Spermacoceae alliances (Figs 1-2) but was sister to the Spermacoceae alliance based on nuclear data [36]. Regarding monophyly of tribes, *Temnopteryx* of the Urophylleae s.l. was sister to all other members of the SCOUT-clade based on nuclear data, whereas Urophylleae s.l. was resolved as monophyletic in the plastome trees (*Temnopteryx* included as sister to the remaining members of the tribe) (Figs 1 and 2). Strongly supported cytonuclear conflict was also found within the tribes Anthospermeae, Argostemmateae, Coussareeae, Lasiantheae, Palicoureeae and Psychotrieae (compare Figs 1 and 2 of the present study with figures 1 and 2 in Thureborn et al. [36]). In passing, it could be noted that the Rubioideae topologies retrieved based on the two analyses of the preferred nuclear dataset in Thureborn et al. [36], i.e., a coalescent-based species tree estimate [figure 1 in reference 36] and a concatenation-based tree estimate [figure 2 in reference 36], differ only concerning one result, the position of Ophiorrhizeae of the SCOUT clade. Here, based on analyses of plastid data, relationships in the SCOUT clade varies among analyses but results lack statistic support in all cases (Figs 1-2 and S1-S2 Figs); there is thus no supported cytonuclear conflict regarding the position of Ophiorrhizeae.

### Simulated plastome trees

Based on simulations under a coalescent model using a nuclear species tree with branch lengths scaled to account for organellar inheritance as a guide tree we generated a distribution of plastome gene trees that would be expected under ILS alone. Support values based on the simulated plastome trees for both the x2 and x4 scalar for the observed plastome relationships are indicated in both the trees inferred from the alignments 1 and 2 (Figs 1 and 2). When using the x4 branch length scalar (to reflect the lower effective population size of dioecious taxa relative to hermaphroditic taxa) support values for the discordant plastome branches were often much lower, as expected as longer branches are associated with lower levels of ILS than are shorter branches. The support for the discordant plastome branches indicates that several empirical plastome relationships were relatively common in simulations, including the branch joining Psychotrieae alliance and Spermacoceae alliance (∼ 29% for both scalars) and the Urophylleae s.l. branch (x2 scalar =7.4%/ x4 scalar = 2.6%), thus in both cases higher than our chosen threshold of 1%. In contrast, some relationships are found in a very small proportion of the simulated trees. For example, the Gaertnereae + Mitchelleae *–* Morindeae clade was found in only 0.9% (x2 scalar) or 0.1% (x4 scalar) of the simulated trees. In one instance, not a single tree contained the observed plastome branch (the x4 scalar trees with the *Durringtonia* + *Opercularia* branch inferred from alignment version 2).

### Basic features of assembled plastomes

We assembled 40 complete plastomes, of which 32 are from the Rubioideae (S1 Table); the remaining plastomes had one to several gaps. The complete circular plastome sequences had a mean of 154 636 bp and ranged in length from 147 802 bp (*Cruckshanksia pumila*) to 158 726 bp (*Ronabea latifolia*). The newly assembled plastome sequences indicated no deviations from the typical quadripartite structure of flowering plants, with a large single copy region (LSC), a small single copy region (SSC) and the inverted repeat region (IR). Also, the typically conserved gene order and gene content of angiosperms were found in most of our newly assembled plastomes. Relative to the conserved angiosperm gene order of the *Coffea arabica* plastome [67] most of the newly obtained plastomes were unrearranged. Important exceptions were the large inversions found in *Carpacoce spermacocea*, *Coprosma longifolia*, *Coprosma rhamnoides*, and *Schismatoclada marojejyensis* (Fig 3). The shortest plastome was that of *Cruckshanksia pumila*. On inspection of whole-plastome alignments, the reduction in size was primarily due to the degradation (e.g., loss and pseudogenization) of the *ndh* gene suite. Otherwise, there were no apparent deviations in gene content compared to the typical gene content of most other angiosperms across the newly assembled plastomes.

**Fig 3.**
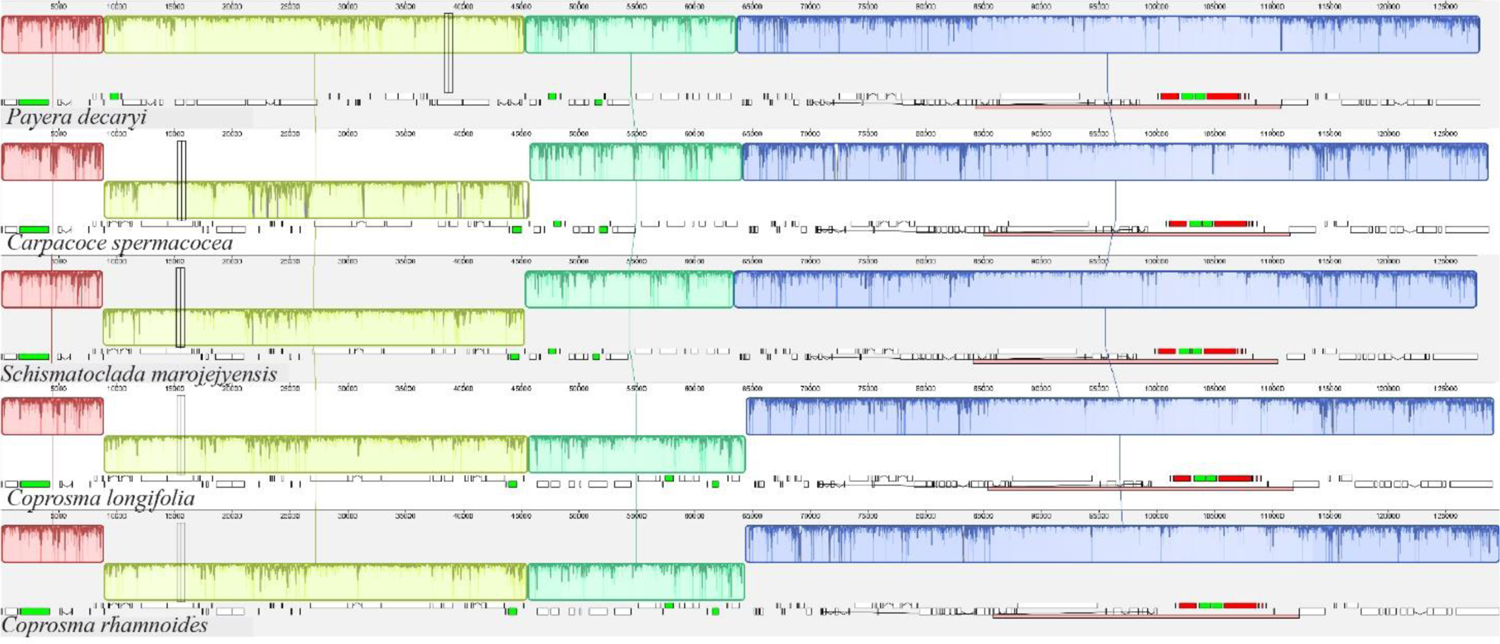
Plastome alignment. The four species with identified rearrangements as well as *Payera decaryi*, which was set as reference (all with one IR only, indicated with pink boxes). Blocks in the same color connected by lines denote locally collinear blocks (LCBs), which represent presumably homologous regions. The LCBs are positioned laterally as they appear in the corresponding plastome. Sunken blocks have an inverted orientation relative to the reference.

## Discussion

### Cytonuclear discordance

Topological conflict between nuclear and organellar phylogenies (cytonuclear discordance) is often detected in phylogenetic studies of angiosperms. Although this phenomenon is frequently detected, investigations of underlying causes have often been hindered by lack of well-resolved species tree estimates. The new sequencing techniques, permitting time- and cost-efficient production of large amounts of genomic data, constitute a possibility to remedy this and increase our understanding of the evolutionary processes that may explain longstanding questions founded in conflicting or unsupported phylogenetic patterns. The relatively few studies that have investigated the source of cytonuclear discordance on deeper phylogenetic levels have considered both hybridization and ILS as likely hypotheses for observed cytonuclear discordance [20, 21, 63, 68–70]. In many studies, neither of these processes has been possible to refute, something that is true also for earlier phylogenomic work on Rubiaceae [35, 37].

Our simulation analyses of plastome data indicate that ILS often is the more likely cause for the observed cytonuclear discordances in the Rubioideae, albeit to varying degrees. Two striking examples of cytonuclear conflict concern the respective placements of Coussareeae and *Temnopteryx*. A sister relationship between Coussareeae and a clade formed by the Psychotrieae and Spermacoceae alliances had long been seen as well-established due to the consistent support for this relationship in phylogenies based on plastid data [e.g., 34, 71-74] as well as mitochondrial data [37]. However, nuclear data [35, 36] support a Coussareeae + Spermacoceae alliance clade, sister to the Psychotrieae alliance. The coalescent-based tree in Thureborn et al. [36] indicated high levels of ILS at the Coussareeae + Spermacoceae alliance branch. In the present study, our analyses show that the discordant plastome tree branch is common in simulations (occurring in c. 29% of the simulated trees) also when accounting for organellar inheritance. This clearly suggests that ILS alone could explain the observed cytonuclear discordance at this part of the tree. In the case involving the placement of *Temnopteryx* the frequencies of simulated plastome trees consistent with the empirical plastome relationship are much lower than those obtained in the case of Coussareeae.

However, the *Temnopteryx +* (remaining) Urophylleae branch is nevertheless relatively common in simulations, in particular under the x2 scalar (x2 scalar = 7.4% / x4 scalar = 2.6%), suggesting that ILS cannot be ruled out and might be the better explanation also for this cytonuclear conflict (rather than plastome introgression).

Another interesting case of cytonuclear discordance concerns the placement of the tribe Gaertnereae of the Psychotrieae alliance. Based on nuclear data [36] Gaertnereae plus the Palicoureeae *–* Psychotrieae clade are sister to a clade, in which Prismatomerideae is sister to Schradereae and Morindeae *–* Mitchelleae. Based on plastome data, Gaertnereae is sister to the Mitchelleae *–* Morindeae clade. The explanation for the observed cytonuclear discordance in this part of the tree may be hybridization followed by chloroplast capture. An ancestor of the Gaertnereae lineage may have captured the plastome from a contemporary ancestor of the Mitchelleae + Morindeae lineage. It should be noted that the observed plastome Gaertnereae + Mitchelleae *–* Morindeae relationship is found among the simulated plastome trees, which makes it difficult to unambiguously reject ILS as explanation for the observed cytonuclear discordance regarding the placement of Gaertnereae, but it is found only in very low numbers. Given the very low fraction (less than 1% under both scalars) of discordant trees detected among the simulated trees, especially under the x4 scalar (0.1% i.e., only 1 of 1000 trees), and the fact that this test can be seen as conservative since at least some gene tree error is expected leading to underestimated branch lengths [31], we argue that (ancient) hybridization and introgression followed by chloroplast capture is the more likely explanation for the discordant position of Gaertnereae.

A complicating factor when considering this hypothetical scenario on the evolutionary history of Gaertnereae is the mitochondrial phylogeny [37], which is in conflict with the plastome phylogeny but completely consistent with the nuclear phylogeny (although Schradereae was not sampled in Rydin et al. [37]). The conflict between mitochondrial and plastome data is surprising as organellar genomes are generally maternally inherited in angiosperms [75]. However, there are documented exceptions to the predominately maternal mode of mitochondrial inheritance [76], for example, the paternal mitochondrial inheritance vs. maternal chloroplast inheritance of *Cucumis* [77] and Pinaceae [78]. Thus, an ancient hybridization scenario may be a possible explanation also regarding the phylogenetic discordance between the two organellar genomes.

The remaining cases of cytonuclear discordance are found within tribes, mainly within the two well-sampled (at the generic level) tribes Argostemmateae and Anthospermeae. The several instances of cytonuclear discordance in those clades are in stark contrast to the lack of observed discordance in the equally fairly well-sampled tribes Ophiorrhizeae and Paederieae. Intergeneric relationships in the Anthospermeae have been problematic to resolve due to conflicting and sometimes poorly supported topologies [79]. Our sampling of this tribe includes eleven of its twelve genera. Only *Nenax* was not sampled since species of this genus have been found intermixed with *Anthospermum* species in a previous study based on Sanger data [79]. The main clades of Anthospermeae [79] are supported in the plastome analyses in the present study, resolving the South African genus *Carpacoce* as sister to a clade divided into an African clade and a Pacific clade. Interpreting the results of our simulations, ILS could well explain most of the observed cytonuclear conflicts in this group, because observed discordant plastome branches are mostly found in a significant fraction among the simulated trees, in most cases in a much higher fraction than our threshold of 1% (Figs 1 and 2).

One exception concerns the discordant *Durringtonia* + *Opercularia* clade found in the plastome tree based on the analysis of alignment 1 (Fig 1). This result, based on alignment 1, is discordant with that based on our plastome alignment 2 (Fig 2), as well as results based on analyses of nuclear genomic data in Thureborn et al. [36]. The clade is not found in any of the simulated plastome trees using the x4 scalar and is rarely found under the x2 scalar (1%).

Similar to the case regarding the position of Gaertnereae this result indicates that ILS alone is an unlikely explanation for the cytonuclear discordance regarding the positions of *Durringtonia* and *Opercularia*; an alternative scenario involving plastid capture following an ancient hybridization event may thus be invoked. It is intriguing that the analysis of our plastome alignment 2 instead resolves *Opercularia* in a clade with *Leptostigma* and *Pomax* (Fig 2), a result concordant with that based on nuclear phylogenies [36, 79]. The conflict is thus not only cytonuclear, but involves a conflicting signal within the plastome as well, something that was indicated also in a previous study based on genome skimming data from a similar sample of Anthospermeae taxa (Thureborn et al. unpublished).

The conflicting topological signals within the Anthospermeae seem to occur across different regions of the plastome (Thureborn et al., unpublished). However, with a concatenated whole plastome framework, the resulting phylogeny depended on the inclusion/exclusion of a specific insertion into an intergenic spacer region between the genes *trnS*^GGA^ and *rps4* in the species of the Pacific clade of Anthospermeae, an insertion of possible mitochondrial origin (Thureborn et al. unpublished). This clade-specific region is not present in alignment version 2 as it was trimmed by the automated gappyout method that removes highly gappy sites.

While the remainder of our inferred plastome relationships in the Rubioideae were robust to the alignment trimming experiment, this result is consistent with the result in Thureborn et al. (unpublished), which showed that small subsets of phylogenomic datasets can alter the topology at specific nodes (see also Shen et al. [80]). A better understanding of the plastome phylogeny and evolution in this particular part of the tree requires further study.

The (mainly) tropical Asian tribe Argostemmateae is here represented by five of the six commonly recognized genera [36, 81–83]. Only *Leptomischus,* which has been placed as sister to the remaining tribe based on *rbcL* data [83] was not sampled. Cytonuclear discordance is extensive in this clade. When comparing our plastome trees (Figs 1 and 2) with the nuclear-based results in Thureborn et al. [36] there is virtually no resemblance between results from the two genomes. As in the Anthospermeae, our simulations indicate that ILS alone might well explain most of this discordance in Argostemmateae because the fraction of discordant trees is high for most nodes (typically much higher than our 1% threshold). An exception is the clade formed by *Argostemma*, *Neohymeopogon* and *Mycetia,* which occurs relatively rarely in simulations (x2 scalar = 1.7% / x4 scalar = 1%) thus indicating that plastome introgression in addition to ILS may be responsible for the observed discord.

The two most comprehensive previous studies focusing on the Argostemmateae were based on a dense taxon sample (although *Clarkella* was not included) and five plastid markers [82] or five plastid markers + nrITS [81]. The sister-relationship between *Mouretia* and an *Argostemma* – *Neohymenopogon – Mycetia* clade found in those studies is congruent with our plastome trees (Figs 1 and 2). The internal phylogeny in this latter clade does, however, differ between the plastome analyses conducted in the present study and that of the previous studies based on a handful of plastid markers. While our analyses support a *Neohymenopogon* + *Mycetia* clade, the two previous studies support an *Argostemma* + *Neohymenopogon* clade (but a single-gene analysis of the plastid *rps16* marker supported the *Neohymenopogon* + *Mycetia* clade [81]). Both these results are thus incongruent with results based on nuclear data [36], and it is interesting to note that the nuclear-based placement of *Mycetia* as sister to *Mouretia* is much more consistent with morphology [81, 84] than either of the plastome-based results are.

### New insights into the phylogeny of Rubioideae as retrieved from analyses of plastid data

#### The deepest splits in the subfamily

Our inference of the Rubioideae phylogeny using whole plastomes (Figs 1 and 2) largely corroborates major relationships found in earlier reconstructions of the evolutionary history of the subfamily using phylogenomic plastome data [35, 38], as well as Sanger-sequenced data of selected plastid markers [34, 73, 85, 86]. There are, however, some noteworthy differences, and some of the differences coincide with areas of the Rubioideae phylogeny that have been notoriously difficult to resolve in the past, for example the deep divergences in the subfamily. Our plastome trees strongly support the SCOUT clade (Seychelleeae, Colletoecemateae, Ophiorrhizeae, and Urophylleae s.l. including *Temnopteryx*) as sister to the rest of the subfamily (Figs 1 and 2). This result was previously established using large amounts of nuclear data [36], but previous studies based on plastid data have not produced results entirely congruent with this result. Instead, these tribes or some of these tribes have been resolved as individual groups successive sisters to the remaining Rubioideae, e.g., Ophiorrhizeae [35, 38] or Colletoecemateae [86] sister to the remaining subfamily. It is possible that these topological differences are due to differences in sampling of characters as well as taxa. Our taxon sampling is much increased compared to previous phylogenomic studies, in that respect more similar to densely sampled Sanger datasets used in the literature. The number of sampled characters is also much increased compared to these earlier studies. Previous phylogenomic studies of plastome data have mainly focused on coding regions; Antonelli et al. [38] analyzed “coding and ribosomal” data assembled from off-target capture sequencing reads, and Wikström et al. [35] analyzed CDS-only and CDS + non-CDS (tRNA rRNA and introns) datasets from near complete and complete plastomes generated via genome skimming reads. In the present study we included information from whole plastomes, including all noncoding regions. Although the generally more conserved coding regions have often been preferred for resolving deeper relationships, fast evolving noncoding regions, such as intergenic regions and introns, have been demonstrated to be highly useful in resolving both deep and shallow relationships in a variety of organismal groups [13, 87–90]. To conclude, it is interesting to note that the SCOUT-clade sister relationship indicated in the nuclear-based species tree in Thureborn et al. [36] has not been reported in previous plastid-based studies. However, with the addition of more data (including non-coding data) the SCOUT-clade relationship is supported also based on plastome data. Similar effects of the inclusion of non-coding data has been observed previously [13].

#### The Psychotrieae alliance

Within the Psychotrieae alliance, our results support Schizocoleeae and Craterispermeae as subsequent sisters to the remaining clade. That clade is in turn split into two major clades (1) a Psychotrieae *–* Palicoureeae clade and (2) a clade where Schradereae and Prismatomerideae are consecutive sisters to a clade, in which Gaertnereae is sister to Morindeae + Mitchelleae. In the plastid Sanger dataset analyzed by Wikström et al. [73] Prismatomerideae was placed outside of the two aforementioned clades. In the plastome-based analyses in Razafimandimbison et al. [74], Wikström et al. [35] and Antonelli et al. [38], Prismatomerideae and Gaertnereae formed a clade sister to sampled members of the Morindeae *–* Mitchelleae clade. Earlier studies have resulted in various topologies (see e.g., the summary in Razafimandimbison et al., [91]), but resolution and/or support were often limited in these studies. As for deep divergences in the subfamily discussed above, differences in sampling (taxa and characters) may be the reasons also for these differing relationships. For example, these previous studies have as a rule sampled only a single representative per tribe, and Schradereae was not included in Wikström et al. [35], and Schradereae and Mitchelleae were not included in Antonelli et al. [38]. Here as in Thureborn et al. [36], we focus on one of the subfamilies of the coffee family and we have sampled the vast majority of the Rubioideae tribes (including all tribes of the Psychotrieae alliance), and we have as a rule sampled more than one representative per tribe.

#### The Spermacoceae alliance

Within the Rubieae complex (here Putorieae *–* Rubieae *–* Theligoneae), Theligoneae is the sister group to Putorieae, which is concordant with previous estimates based on nuclear data [36], and with the plastome-based tree of Antonelli et al. [38] and the Sanger plastid tree of Yang et al. [92]. Plastid-based results in other studies have found Theligoneae as sister to Rubieae, i.e., the plastome trees in Wikström et al. [35] and results based on Sanger-sequenced data [34, 71, 93–95]. It is interesting to note that the two previous phylogenomic studies that both mainly relied on CDS plastome data and have similar taxon sampling density (i.e., usually one representative per tribe) show strongly conflicting topologies. This may possibly indicate that the specific species chosen to represent each tribe (which differ among these studies) are of importance for topological results in this case.

A surprising result in Antonelli et al. [38] was the placement of *Cyanoneuron pedunculatum* of the tribe Cyanoneuroneae, which was nested in the Psychotrieae alliance in their plastome tree. We used the same raw data of *Cyanoneuron pedunculatum* in the present study, and in addition newly produced data from another species of *Cyanoneuron, C. cyaneum.* The resulting topology (Figs 1 and 2) agrees with those of other previous studies based on plastid data as well as nuclear data [36, 38, 79, 82]: *Cyanoneuron pedunculatum* and *C. cyaneum* are sisters and included in the Spermacoceae alliance. It is likely that the highly unexpected position of Cyanoneuroneae/*Cyanoneuron pedunculatum* in the Psychotrieae alliance in Antonelli et al. [38] is an artefact. For example, while our plastome of *Cyanoneuron cyaneum* is nearly complete, the data from the *Cyanoneuron pedunculatum* sample has the lowest plastome sequence recovery of all samples included in the present study. The addition of a more complete sequence from the Cyanoneuroneae may thus have been helpful for resolving the phylogenetic position of the tribe. Furthermore, a denser taxonomic sampling may also have improved accuracy of results. In addition to an additional sample of Cyanoneuroneae (*Cyanoneuron cyaneum*), we also included representation of the closely related tribe Dunnieae.

### On data sampling and methodological choices in phylogenetic studies

The plastome is typically considered a single linkage group, uniparentally (usually maternally) inherited and often treated as a single “supergene” in phylogenetic analyses. This traditional treatment of plastome data in phylogenetic studies has recently become subject to some debate [4, 96]. However, assuming that the entire plastome indeed share the same phylogenetic history, reasons for phylogenetic conflict within the plastome should primarily be sought among analytical factors such as sampling strategy, rather than biological causes. A dense taxon sampling is of high importance for accurate estimations of tree topology [97, 98], and is when combined with additional sampling of characters often considered a remedy for resolving long-standing phylogenetic questions [99]. Among other things, a denser taxon sampling improves estimation of model parameters [97, 100, 101], and studies have shown that increased taxon sampling density is more important for accuracy of phylogenetic estimation than model complexity [17, 102].

Data recoding [103] is, in addition to increased sampling and model complexity, another strategy that can improve phylogenetic accuracy. Wikström et al. [35] applied degenerate recoding to their CDS data, a method that uses ambiguity codes to remove potentially misleading synonymous substitutions caused by saturation or compositional bias [104]. While this treatment supported an alternative Cinchonoideae topology, the Rubioideae phylogeny was unaffected [35]. Nevertheless, we wanted to explore the impact of potential saturation and compositional bias on our data using RY-coding, a method that is applicable to all kinds of nucleotide sequence data, not only to CDS data. RY-coding normalizes the four nucleotide states by recoding them into two states (purines and pyrimidines). The use of ambiguity codes does, however, come at the expense of phylogenetic information [56, 58]. Here, RY-coding only affected one poorly supported ingroup branch and reduced support for some, and our results suggest that our estimates of the Rubioideae phylogeny are robust to potential artifacts due to base composition or saturation. However, this does not exclude the possibility that individual loci may be affected by saturation and compositional bias although if so, not influential enough to alter results of analyses of the concatenated dataset.

Several studies have investigated the impact of data partitioning, and it has been shown that topological results retrieved from unpartitioned vs. partitioned plastome data are identical or almost identical [89, 105, 106]. Furthermore, a recent study [107] showed that it was mainly nodes with poor support that were affected by the choice of partitioning scheme. When datasets increased in size the dependency on partitioning scheme decreased [107], again highlighting the importance of sampling. Thus, while the choices of data partitioning have varied between studies (e.g., unpartitioned ML analyses in the present study, partitioned ML analyses in Antonelli et al. [38] and Wikström et al. [35]), this seems unlikely as explanatory factor for the topological results that differ between these studies.

### Plastome evolution in the Rubioideae

The successful recovery of large amounts of plastome data from off-target reads employed in the present study can probably be attributed to the generality of the probe set we used in Thureborn et al. [36]. The target capture sequencing reads in Thureborn et al. [36] were captured using the universal Angiosperms353 probe kit [24], and the on-target read fraction was on average 15.8% [36]. Had we used a more specific probe set specifically designed for the Rubioideae, thus with a higher enrichment efficiency, additional sequencing effort would possibly have been necessary to reach sufficient amount of reads for plastome assembly. Our main focus in the present study was to address phylogenetic questions and phylogenetic discordance within the Rubioideae, however, the *de novo* and combined reference-guided + *de novo* approach we used for plastome assembly facilitates a brief discussion on plastome evolution in the group.

With few exceptions, the here newly obtained plastomes are highly similar to the ancestral angiosperm plastome in size, structure, gene content, and gene order [108]. This corroborates the overall conserved nature of plastomes that has been reported for the Rubiaceae in comparative studies focusing on subfamily Ixoroideae [109], the *Coffea* alliance (Ixoroideae) [110] and *Leptodermis* (Rubioideae) [111]. However, Thureborn et al. (unpublished data) found two large inversions (36-kb and 19-kb) in *Coprosma rotundifolia* in a phylogenomic study of the tribe Anthospermeae. The 36-kb inversion has previously been described for the genistoid clade of the Fabaceae by Martin et al. [112] who predicted that it may occur in additional groups as well since it is likely mediated by a 29-bp inverted repeats located in the 3’-ends of the *trnS*^GGA^ and *trnS*^GCU^ genes. The *trnS*^GGA^ *– trnS*^GCU^ inversion has, however, rarely been reported since although it has been found in distantly related genera of the Fabaceae [113, 114] and thus recently also in one representative of the genus *Coprosma* of the Rubiaceae (Thureborn et al. unpublished data).

Here we found the 36-kb and the 19-kb inversions in both the newly assembled *Coprosma* species. The 36-kb inversion was also identified in the newly assembled plastomes of *Carpacoce spermacocea* (Anthospermeae) and in *Schismatoclada marojejyensis* (Danaideae). Interestingly, another sample of *Carpacoce spermacocea* has previously been sequenced (Thureborn et al., unpublished data), and it did not have the 36-kb insertion. Alternative arrangements can thus be found even within the same species. Homoplastic inversions have also been found in other angiosperm lineages [115–117], including intraspecific differences [118], indicating that large inversion events generally should be treated cautiously when used as phylogenetic characters. However, so far, the presence of both the 36-kb and 19-kb inversions found in the three species of *Coprosma* (the present study and Thureborn et al., unpublished data) indicates that it is a potential synapomorphy for this genus.

Transfer of mitochondrial DNA into the plastome is regarded as a rare event, although it has been reported [119, 120]. An insertion of putative mitochondrial DNA into the plastome of the members of the Pacific clade of Anthospermeae was discovered recently (Thureborn et al., unpublished data) and is a possible synapomorphy for that clade. The here newly sequenced plastomes of the members of the Pacific clade have sequences that are positionally homologous to those with the previously identified insertions, reinforcing that the putative mitochondrial insertion is a synapomorphy for this clade. It should be noted that the putative mitochondrial insert is found in the *trnS*^GGA^ – *rps4* IGS in six of the seven genera of the Pacific clade, but in *Coprosma*, due to the 19-kb inversion, sequences homologous to the *trnS*^GGA^ – *rps4* IGS (including putative mitochondrial sequence) is located in the *trnS*^GCU^ *– petA* and *rps4 – psbJ* IGSs of the *Coprosma* plastomes.

The here observed *ndh* gene suite degradation (including loss and pseudogenization) in *Cruckshanksia pumila* is the first report of degradation of this gene suite in Rubiaceae plastomes. Although rather rare, the degradation of the *ndh* genes has been reported in several other land plants. *Cruckshanksia pumila* is a (typically) annual species living in the northern arid regions of Chile [121]. Degradation of plastid genes is common in parasitic and mycoheterotrophic plants, in which the need for photosynthesis has become redundant [122, 123], but degradation of *ndh* genes has also been reported in a diverse set of autotrophic plants, including gymnosperms [124, 125] and angiosperms. In angiosperms *ndh* genes have been lost in carnivorous plants [126, 127], and aquatic plants [128] but also in other distantly related autotrophs such as the alpine ranunculalean herb *Kingdonia uniflora* [129], *Gentiana* (Gentianales) [130], *Erodium* (Geraniales) [131], orchids [132, 133], and cacti where the loss of *ndh* is correlated with CAM photosynthesis [134]. The *ndh* complex is suggested to have an important role in improving photosynthesis under abiotic stress conditions, but may be superfluous under mild environmental conditions [135].

## Acknowledgements

We thank Bodil Cronholm, Martin Irestedt (the Swedish Museum of Natural History), and Anbar Khodabandeh (the Royal Swedish Academy of Sciences) for technical assistance, and we acknowledge support from the National Genomics Infrastructure in Stockholm (Science for Life Laboratory, the Knut and Alice Wallenberg Foundation and the Swedish Research Council), and SNIC (Uppsala Multidisciplinary Center for Advanced Computational Science) for assistance with massively parallel sequencing and access to the UPPMAX computational infrastructure.

## Data accessibility statement

Newly generated plastome sequences have been deposited in the Dryad Digital Repository: https://doi.org/10.5061/dryad.mpg4f4r67, and the previously published plastome sequences as well as raw sequencing reads are available in the GenBank and ENA repositories (for accessions, see S1 Table).

## Author contributions

**Conceptualization**: Olle Thureborn, Catarina Rydin, Niklas Wikström, Sylvain G. Razafimandimbison

**Data curation**: Olle Thureborn

**Formal analysis**: Olle Thureborn

**Funding acquisition**: Catarina Rydin

**Investigation**: Olle Thureborn

**Project administration**: Catarina Rydin

**Resources**: Catarina Rydin

**Supervision**: Catarina Rydin, Niklas Wikström, Sylvain G. Razafimandimbison

**Validation**: Olle Thureborn, Catarina Rydin, Niklas Wikström, Sylvain G. Razafimandimbison

**Visualization**: Olle Thureborn, Catarina Rydin

**Writing – original draft**: Olle Thureborn

**Writing – review & editing**: Olle Thureborn, Catarina Rydin, Niklas Wikström, Sylvain G. Razafimandimbison

## Supporting information

**S1 Table.** Taxa, voucher/source, ENA/GenBank, assembly, and sequencing information for sequences used in this study. (pdf)

**S1 Fig.** Plastome phylogeny inferred from the maximum likelihood analysis of the RY-coded version of the alignment 1. (pdf)

**S2 Fig.** Plastome phylogeny inferred from the maximum likelihood analysis of the RY-coded version of the alignment 2. (pdf)

