## Supplementary Material for "Plastid phylogenomics and cytonuclear discordance in Rubioideae, Rubiaceae"

**S1 Table. Taxa, voucher/source, ENA/GenBank, assembly, and sequencing information for sequences used in this study.**

Taxa with fully assembled plastomes are indicated in grey.

| Lab ID | Species | Family (Subfamily) | Tribe (Rubiaceae only) | Collection/Source | Year | Collection locality | Assembly strategy* |
|---|---|---|---|---|---|---|---|
| | Alstonia scholaris (L.) R.Br. | Apocynaceae | | Antonelli et al. (2021)/PAFTOL | | | RG-only |
| | Chironia baccifera L. | Gentianaceae | | Antonelli et al. (2021)/PAFTOL | | | RG-only |
| | Geniostoma borbonicum (Lam.) Spreng. | Loganiaceae | | Antonelli et al. (2021)/PAFTOL | | | RG+de novo |
| BD003 | Acranthera grandiflora Bedd. | Rubiaceae | Coptosapelteae | J. Klackenberg & R. Lundin 541 (S) | 1982 | India, Tamil Nadu | RG+de novo |
| AY008 | Coptosapelta diffusa (Champ. ex Benth.) Steenis | Rubiaceae | Coptosapelteae | Steward et al 594 (S) | 1931 | China | RG+de novo |
| AZ040 | Luculia pinceana Hook. | Rubiaceae | Luculieae | Thin et al. 3061 (AAU) | | | de novo |
| DE068 | Cinchona calisaya Wedd. | Rubiaceae (Cinchonoideae) | Cinchoneae | Razafimandim. & Razafimanant. 471 (UPS) | 2002 | Madagascar | RG+de novo |
| CC071 | Deppea grandiflora Schltdl. | Rubiaceae (Cinchonoideae) | Hamelieae | Marino Rosas R. 1015 (P) | | | RG-only |
| DE066 | Paracorynanthe antankarana Capuron ex J.-F.Leroy | Rubiaceae (Cinchonoideae) | Hymenodictyeae | B. Bremer et al. 5156 (S) | 2008 | Madagascar | RG+de novo |
| | Corynanthe pachyceras K.Schum. | Rubiaceae (Cinchonoideae) | Naucleeae | Antonelli et al. (2021)/PAFTOL | | | RG-only |
| AG098 | Rondeletia intermixta Britton | Rubiaceae (Cinchonoideae) | Rondeletieae | Rova et al. 2245 (GB) | 1995 | Cuba | RG+de novo |
| AP035 | Airosperma vanuense S.P.Darwin | Rubiaceae (Ixoroideae) | Airospermeae | Smith 8214 (P) | 1953 | Fiji | RG+de novo |
| | Razafimandimbisonia minor (Baill.) Kainul. & B.Bremer | Rubiaceae (Ixoroideae) | Alberteae | Antonelli et al. (2021)/PAFTOL | | | RG-only |
| BE034 | Wendlandia dasythyrsa Miq. | Rubiaceae (Ixoroideae) | Augusteae | Christensen, H. M. 460 (AAU) | 1993 | Malaysia | RG+de novo |
| BD032 | Alseis lugonis L.Andersson | Rubiaceae (Ixoroideae) | Condamineeae | Bremer et al. 3353 (UPS) | 1995 | Ecuador | RG-only |
| | Emmenopterys henryi Oliv. | Rubiaceae (Ixoroideae) | Condamineeae | Antonelli et al. (2021)/PAFTOL | | | RG-only |
| CX014 | Ixora javanica (Blume) DC. | Rubiaceae (Ixoroideae) | Ixoreae | Puff 000512-1/2 (WU) | | Vietnam | RG+de novo |
| | Jackiopsis ornata (Wall.) Ridsdale | Rubiaceae (Ixoroideae) | Jackieae | Antonelli et al. (2021)/PAFTOL | | | RG-only |
| | Heinsia crinita (Wennberg) G.Taylor | Rubiaceae (Ixoroideae) | Mussaendeae | Antonelli et al. (2021)/PAFTOL | | | RG-only |
| AV015 | Molopanthera paniculata Turcz. | Rubiaceae (Ixoroideae) | Posoquerieae | Williams & Assis 6861 (S) | 1945 | Brazil | de novo |
| CG080 | Sabicea marojejyensis Razafim. & J.S.Mill. | Rubiaceae (Ixoroideae) | Sabiceeae | Bremer et al. 5293 (S) | 2008 | Madagascar | de novo |
| | Cyclophyllum barbatum (G.Forst.) N.Hallé & J.Florence | Rubiaceae (Ixoroideae) | Vanguerieae | Antonelli et al. (2021)/PAFTOL | | | RG+de novo |
| CA098 | Glionnetia sericea (Baker) Tirveng. | Rubiaceae (Ixoroideae) | | Beaver 3 (S) | 2009 | Seychelles | de novo |
| DE070 | Anthospermum thymoides Baker | Rubiaceae (Rubioideae) | Anthospermeae | Thureborn et al. 33 (S) | 2017 | Madagascar | RG+de novo |
| DE067 | Carpacoce spermacocea (Rchb. ex Spreng.) Sond. | Rubiaceae (Rubioideae) | Anthospermeae | Bremer et al. 4385 (UPS) | 2002 | South Africa | RG+de novo |
| | Coprosma longifolia A.Gray | Rubiaceae (Rubioideae) | Anthospermeae | Antonelli et al. (2021)/PAFTOL | | | RG+de novo |
| CY012 | Coprosma rhamnoides A.Cunn. | Rubiaceae (Rubioideae) | Anthospermeae | Tibell NZ46 (UPS) | 1980 | New Zealand | RG+de novo |
| DE081 | Durringtonia paludosa R.J.F.Hend. & Guymer | Rubiaceae (Rubioideae) | Anthospermeae | Thompson 18m11 (BRI) | 2001 | Australia, Queensland | RG+de novo |
| CX098 | Galopina aspera (Eckl. & Zeyh.) Walp. | Rubiaceae (Rubioideae) | Anthospermeae | Phillipson 1461 (UPS) | 1986 | South Africa | RG+de novo |
| CX099 | Leptostigma pilosum (Benth.) Fosberg | Rubiaceae (Rubioideae) | Anthospermeae | Asplund 7171 (UPS) | 1939 | Ecuador | RG+de novo |
| CY003 | Nertera dichondrifolia (A.Cunn.) Hook.f. | Rubiaceae (Rubioideae) | Anthospermeae | Tibell NZ119 (UPS) | 1980 | New Zealand | de novo |
| DE075 | Normandia neocaledonica Hook.f. | Rubiaceae (Rubioideae) | Anthospermeae | Selling 125b (S) | 1949 | New Caledonia | RG+de novo |
| CY089 | Opercularia volubilis R.Br. ex Benth. | Rubiaceae (Rubioideae) | Anthospermeae | B.J. Lepschi & B.A. Fuhrer BJL 3671 (P) | 1997 | Australia | de novo |
| CY051 | Phyllis nobla L. | Rubiaceae (Rubioideae) | Anthospermeae | Wikström et al. 76 (S) | 1994 | Tenerife | de novo |
| DE074 | Pomax umbellata (Gaertn.) Sol. ex A.Rich. | Rubiaceae (Rubioideae) | Anthospermeae | Halford Q9744 (BRI) | 2009 | Australia | RG+de novo |
| BL040 | Argostemma elatostemma Hook.f. | Rubiaceae (Rubioideae) | Argostemmateae | Bremer and Bremer 1722 (S) | 1979 | Malaysia, Sarawak | RG+de novo |
| BU095 | Clarkella nana (Edgew.) Hook.f. | Rubiaceae (Rubioideae) | Argostemmateae | Maxwell 02-252 (MO) | | | RG-only |
| CH091 | Mouretia larsenii Tange | Rubiaceae (Rubioideae) | Argostemmateae | van Beusekom 4743 (P) | 1972 | Thailand | de novo |
| CD007 | Mycetia bracteata Hutch. | Rubiaceae (Rubioideae) | Argostemmateae | Steward & Cheo 1105 (S) | 1933 | China | de novo |
| | Mycetia Reinw. | Rubiaceae (Rubioideae) | Argostemmateae | Antonelli et al. (2021)/PAFTOL | | | RG-only |
| BK078 | Neohymenopogon parasiticus (Wall.) Bennet | Rubiaceae (Rubioideae) | Argostemmateae | Bremer 2743 (UPS) | | Cult. | RG+de novo |
| BD002 | Colletoecema dewevrei (De Wild.) E.M.A.Petit | Rubiaceae (Rubioideae) | Colletoecemateae | Lisowski 47195 (K) | 1977 | Congo | NA |
| BA061 | Coccocypselum condalia Pers. | Rubiaceae (Rubioideae) | Coussareeae | Persson, C. & Gustafsson, C. 246 (GB) | 1996 | Bolivia | RG+de novo |

**S1 Table. Taxa, voucher/source, ENA/GenBank, assembly, and sequencing information for sequences used in this study.**

Taxa with fully assembled plastomes are indicated in grey.

| Lab ID | Species | Family (Subfamily) | Tribe (Rubiaceae only) | Collection/Source | Year | Collection locality | Assembly strategy* |
|---|---|---|---|---|---|---|---|
| DA042 | Cruckshanksia pumila Clos in C.Gay | Rubiaceae (Rubioideae) | Coussareeae | Taylor et al. 10679 (MO) | 1991 | Chile | RG+de novo |
| DB007 | Faramea multiflora A.Rich. ex DC. | Rubiaceae (Rubioideae) | Coussareeae | Salino 3825 (MO ) | 1997 | Brazil | RG+de novo |
| | Craterispermum Benth. 2 | Rubiaceae (Rubioideae) | Craterispermeae | Antonelli et al. (2021)/PAFTOL | | | RG-only |
| CK013 | Craterispermum Benth. 1 | Rubiaceae (Rubioideae) | Craterispermeae | Razafimandimbison et al. 1168 (S) | 2011 | Madagascar | RG+de novo |
| CL038 | Craterispermum schweinfurthii Hiern | Rubiaceae (Rubioideae) | Craterispermeae | J. D & E. G. Chapman 9364 (UPS) | 1988 | Malawi | RG+de novo |
| BZ084 | Cyanoneuron cyaneum (Hallier f.) Tange | Rubiaceae (Rubioideae) | Cyanoneuroneae | Bogner 1457 (L) | | Borneo, Indonesia | RG+de novo |
| | Cyanoneuron pedunculatum Tange | Rubiaceae (Rubioideae) | Cyanoneuroneae | Antonelli et al. (2021)/PAFTOL | | | RG-only |
| DB070 | Danais nigra Homolle | Rubiaceae (Rubioideae) | Danaideae | N. S. Rasoanaivo & A. J. Tahinarivony 69 (S) | 2012 | Madagascar | RG+de novo |
| DB077 | Payera decaryi (Homolle) Buchner & Puff | Rubiaceae (Rubioideae) | Danaideae | Krüger & Razafimandimbison 74 (S) | 2010 | Madagascar | de novo |
| DC094 | Schismatoclada marojejyensis Humbert | Rubiaceae (Rubioideae) | Danaideae | Bremer et al. 5309 (S) | 2008 | Madagascar | de novo |
| BC005 | Dunnia sinensis Tutcher | Rubiaceae (Rubioideae) | Dunnieae | Xinhui 16 (in Ge et al, 2002) | | | RG+de novo |
| CI009 | Gaertnera obovata Baker | Rubiaceae (Rubioideae) | Gaertnereae | Razafimandimbison et al. 980 (S) | 2011 | Madagascar | RG+de novo |
| | Gaertnera rotundifolia Bojer | Rubiaceae (Rubioideae) | Gaertnereae | Antonelli et al. (2021)/PAFTOL | | | RG-only |
| CA030 | Pagamea capitata Benth. | Rubiaceae (Rubioideae) | Gaertnereae | Pipoly 9176 (MEXU) | 1986 | Guyana | RG+de novo |
| AP063 | Otiophora caerulea (Hiern) Bullock | Rubiaceae (Rubioideae) | Knoxieae | Dessein 367 (BR) | 2004 | Zambia | RG+de novo |
| AI014 | Chamaepentas hindsioides (K.Schum.) Kårehed & B.Bremer | Rubiaceae (Rubioideae) | Knoxieae | Iversen et al. 85101 (UPS) | 1985 | Tanzania | de novo |
| DE064 | Triainolepis xerophila (Bremek.) Kårehed & B.Bremer | Rubiaceae (Rubioideae) | Knoxieae | Thureborn et al. 11 (S) | 2017 | Madagascar | RG+de novo |
| CQ058 | Lasianthus Jack | Rubiaceae (Rubioideae) | Lasiantheae | Razafimandimbison et al. 718 (S) | 2009 | Vietnam | RG-only |
| CH072 | Lasianthus strigosus Wight | Rubiaceae (Rubioideae) | Lasiantheae | Bremer & Bremer 3902 (UPS) | 1998 | Australia | RG+de novo |
| AZ001 | Ronabea latifolia Aubl. | Rubiaceae (Rubioideae) | Lasiantheae | Contreras 9152 (S) | | Guatemala | RG+de novo |
| | Saldinia aegialodes Bremek. | Rubiaceae (Rubioideae) | Lasiantheae | Antonelli et al. (2021)/PAFTOL | | | RG-only |
| AA052 | Saldinia pallida Bremek. | Rubiaceae (Rubioideae) | Lasiantheae | Bremer et al. 4038-BB38 (UPS) | 2000 | Madagascar | RG+de novo |
| BE046 | Trichostachys microcarpa K.Schum. | Rubiaceae (Rubioideae) | Lasiantheae | Masens 834 (BR) | 1991 | Congo | RG+de novo |
| BZ092 | Mitchella repens L. | Rubiaceae (Rubioideae) | Mitchelleae | Atha and Gonzalez 1443a (MEXU) | 1997 | United States, Texas | NA |
| AQ075 | Appunia guatemalensis Donn.Sm. | Rubiaceae (Rubioideae) | Morindeae | Contreras 8983 (S) | 1969 | Guatemala | RG+de novo |
| | Coelospermum paniculatum F.Muell. | Rubiaceae (Rubioideae) | Morindeae | Antonelli et al. (2021)/PAFTOL | | | RG-only |
| BZ099 | Gynochthodes officinalis (F.C.How) Razafim. & B.Bremer | Rubiaceae (Rubioideae) | Morindeae | Krüger et al. 9 (S) | 2009 | Vietnam | RG+de novo |
| CV018 | Morinda citrifolia L. | Rubiaceae (Rubioideae) | Morindeae | Razafimandimbison 1212a (S) | 2013 | Madagascar | de novo |
| AX046 | Lerchea bracteata Valeton | Rubiaceae (Rubioideae) | Ophiorrhizeae | Axelius 343 (S) | 1983 | Indonesia, Sumatra | RG+de novo |
| | Kajewskiella trichantha Merr. & L.M.Perry | Rubiaceae (Rubioideae) | Ophiorrhizeae | Antonelli et al. (2021)/PAFTOL | | | RG-only |
| CH086 | Neurocalyx zeylanicus Hook. | Rubiaceae (Rubioideae) | Ophiorrhizeae | Bremer & Bremer 937 (S) | 1977 | Sri Lanka | RG+de novo |
| CY100 | Ophiorrhiza darwinii Razafim. & Rydin | Rubiaceae (Rubioideae) | Ophiorrhizeae | Swenson et al. 1411 (S) | 2013 | Vietnam | RG+de novo |
| CZ012 | Ophiorrhiza mungos L. | Rubiaceae (Rubioideae) | Ophiorrhizeae | USA Typhus Commission 525 (S) | | | RG+de novo |
| | Ophiorrhiza winkleri Valeton | Rubiaceae (Rubioideae) | Ophiorrhizeae | Antonelli et al. (2021)/PAFTOL | | | RG-only |
| | Paederia thouarsiana Baill. | Rubiaceae (Rubioideae) | Paederieae | Antonelli et al. (2021)/PAFTOL | | | RG+de novo |
| P0085 | Leptodermis potaninii Batalin | Rubiaceae (Rubioideae) | Paederieae | Andreasen 230 (UPS) | 1993 | Cult. Quarryhill Bot. Gard. | RG+de novo |
| CA022 | Paederia ciliata (Bartl. ex DC.) Standl. | Rubiaceae (Rubioideae) | Paederieae | Campos et al. 4919 (MEXU) | 1992 | Mexico | RG+de novo |
| BX093 | Pseudopyxis heterophylla (Miq.) Maxim. | Rubiaceae (Rubioideae) | Paederieae | K.Å. Dahlstrand s.n. (GB) | 1950 | Japan | RG+de novo |
| C0005 | Serissa foetida (L.f.) Lam. | Rubiaceae (Rubioideae) | Paederieae | Bremer 2735 (UPS) | 1988 | Cult. in Kew Bot. Gard. | RG+de novo |
| B0110 | Spermadictyon suaveolens Roxb. | Rubiaceae (Rubioideae) | Paederieae | Bremer 3133 (UPS) | 1988 | Cult. in Paris Bot. Gard. | RG+de novo |
| AE034 | Rudgea recurva Müll.Arg. | Rubiaceae (Rubioideae) | Palicoureeae | Pirani et al. 4899 (SPF) | 2001 | Brazil | RG+de novo |
| BH070 | Palicourea alpina (Sw.) DC. | Rubiaceae (Rubioideae) | Palicoureeae | Rova 2246 (GB) | 1995 | Cuba | RG+de novo |
| | Palicourea nitidella (Müll.Arg.) Standl. | Rubiaceae (Rubioideae) | Palicoureeae | Antonelli et al. (2021)/PAFTOL | | | RG-only |

**S1 Table. Taxa, voucher/source, ENA/GenBank, assembly, and sequencing information for sequences used in this study.**

Taxa with fully assembled plastomes are indicated in grey.

| Lab ID | Species | Family (Subfamily) | Tribe (Rubiaceae only) | Collection/Source | Year | Collection locality | Assembly strategy* |
|---|---|---|---|---|---|---|---|
| CM050 | Puffia gerrardii (Baker) Razafim. & B.Bremer | Rubiaceae (Rubioideae) | Palicoureeae | Razafimandimbison et al. 1244 (S) | 2013 | Madagascar | RG+de novo |
| | Perama dichotoma Poepp. | Rubiaceae (Rubioideae) | Perameae | Antonelli et al. (2021)/PAFTOL | | | RG-only |
| AM028 | Perama hirsuta Aubl. | Rubiaceae (Rubioideae) | Perameae | Andersson et al. 1990 (GB) | | French Guiana | de novo |
| CA031 | Prismatomeris fragrans E.T.Geddes | Rubiaceae (Rubioideae) | Prismatomerideae | Kainulainen et al. 39 (S) | 2009 | Vietnam | NA |
| | Prismatomeris Thwaites | Rubiaceae (Rubioideae) | Prismatomerideae | Antonelli et al. (2021)/PAFTOL | | | RG-only |
| CB078 | Rennellia subsessilis (King & Gamble) Razafim. & Rydin | Rubiaceae (Rubioideae) | Prismatomerideae | Y.W. Low & Wong LYW 359 (KLU) | | Malaysia | RG+de novo |
| AF076 | Psychotria ankarensis (Bremek.) Razafim. & B.Bremer | Rubiaceae (Rubioideae) | Psychotrieae | Razafimandimbison et al. 405 (UPS) | 2002 | Madagascar | de novo |
| AG024 | Psychotria mahonii C.H.Wright | Rubiaceae (Rubioideae) | Psychotrieae | Luke 8370 (UPS) | 2002 | Kenya | RG-only |
| CL028 | Calycosia lageniformis (Gillespie) A.C.Sm. | Rubiaceae (Rubioideae) | Psychotrieae | Callmander et al. 962 (S) | | Fiji | RG+de novo |
| | Calycosia petiolata A.Gray | Rubiaceae (Rubioideae) | Psychotrieae | Antonelli et al. (2021)/PAFTOL | | | RG-only |
| | Chaetostachydium barbatum Ridsdale | Rubiaceae (Rubioideae) | Psychotrieae | Antonelli et al. (2021)/PAFTOL | | | RG-only |
| | Dolianthus montiswilhelmii (P.Royen) A.P.Davis | Rubiaceae (Rubioideae) | Psychotrieae | Antonelli et al. (2021)/PAFTOL | | | RG+de novo |
| | Psychotria pandurata Verdc. | Rubiaceae (Rubioideae) | Psychotrieae | Antonelli et al. (2021)/PAFTOL | | | RG-only |
| | Plocama calabrica (L.f.) M.Backlund & Thulin | Rubiaceae (Rubioideae) | Putorieae | Antonelli et al. (2021)/PAFTOL | | | RG-only |
| AH008 | Plocama tinctoria (Balf.f.) M.Backlund & Thulin | Rubiaceae (Rubioideae) | Putorieae | Thulin 10946 (UPS) | 2002 | Somalia | RG+de novo |
| AH009 | Plocama dubia (Aitch. & Hemsl.) N. Backlund & Thulin | Rubiaceae (Rubioideae) | Putorieae | Rafei & Zangooei 25651 (FUHM) | | | RG+de novo |
| | Rubia peregrina L. | Rubiaceae (Rubioideae) | Rubieae | Antonelli et al. (2021)/PAFTOL | | | RG-only |
| AG064 | Sherardia arvensis L. | Rubiaceae (Rubioideae) | Rubieae | Andreasen 345 (SBT) | 2002 | Cult. in Bergius Bot. Gard. | RG+de novo |
| M0004 | Didymaea alsinoides (Schltdl. & Cham.) Standl. | Rubiaceae (Rubioideae) | Rubieae | Keller 1901 (CAS) | | | RG+de novo |
| DE065 | Galium polyacanthum (Baker) Puff | Rubiaceae (Rubioideae) | Rubieae | Thureborn et al. 32 (S) | 2017 | Madagascar | RG+de novo |
| T0054 | Kelloggia galioides Torr. | Rubiaceae (Rubioideae) | Rubieae | Holmgren et al. 2437 (UPS) | 1965 | United States, Utah | de novo |
| BO005 | Rubia cordifolia subsp. conotricha (Gand.) Verdc. | Rubiaceae (Rubioideae) | Rubieae | P.A. Luke & W.R.Q Luke 9510 (UPS) | 2003 | Tanzania | RG+de novo |
| CH081 | Schizocolea linderi (Hutch. & Dalziel) Bremek. | Rubiaceae (Rubioideae) | Schizocoleeae | Adam 20116 (UPS) | 1964 | Liberia | NA |
| BZ091 | Lecananthus erubescens Jack | Rubiaceae (Rubioideae) | Schradereae | Van Niel 3406 (L) | | Borneo, Indonesia | RG+de novo |
| CX040 | Schradera nervulosa (Stapf) Puff, R.Buchner & Greimler | Rubiaceae (Rubioideae) | Schradereae | Puff 961216-2/2 (WU) | | Borneo (Sabah) | RG+de novo |
| CA038 | Schradera rotundata Standl. ex Steyerm. | Rubiaceae (Rubioideae) | Schradereae | Gentry 5641 (S) | | Panama | de novo |
| CM080 | Seychellea sechellarum (Baker) Razafim., Kainul. & Rydin | Rubiaceae (Rubioideae) | Seychelleeae | C. Morel 57a (SEY) | | Seychelles | RG+de novo |
| CG025 | Diodella sarmentosa (Sw.) Bacigalupo & E. L. Cabral | Rubiaceae (Rubioideae) | Spermacoceae | Bremer et al. 5229 (S) | 2008 | Madagascar | RG+de novo |
| BZ040 | Exallage chrysotricha (Palib.) Neupane & N.Wikstr. | Rubiaceae (Rubioideae) | Spermacoceae | Lin Qinzhong 2004180 (MO) | 2004 | China | RG+de novo |
| DE079 | Oldenlandia herbacea (L.) Roxb. | Rubiaceae (Rubioideae) | Spermacoceae | Thureborn et al. 27 (S) | 2017 | Madagacar | de novo |
| | Spermacoce L. | Rubiaceae (Rubioideae) | Spermacoceae | Antonelli et al. (2021)/PAFTOL | | | RG-only |
| | Theligonum cynocrambe L. | Rubiaceae (Rubioideae) | Theligoneae | Antonelli et al. (2021)/PAFTOL | | | RG+de novo |
| BV076 | Theligonum japonicum Ôkubo & Makino | Rubiaceae (Rubioideae) | Theligoneae | Togashi 6807 (P) | 1968 | Japan | RG+de novo |
| AX027 | Amphidasya longicalycina (Dwyer) C.M.Taylor | Rubiaceae (Rubioideae) | Urophylleae | Huber 2963 (CR) | | Costa Rica | RG+de novo |
| AS084 | Temnopteryx sericea Hook.f. | Rubiaceae (Rubioideae) | Urophylleae | Tabak 999 (WAG) | | Gabon | RG+de novo |
| | Urophyllum cyphandrum Stapf | Rubiaceae (Rubioideae) | Urophylleae | Antonelli et al. (2021)/PAFTOL | | | RG-only |
| AY100 | Raritebe palicoureoides Wernham | Rubiaceae (Rubioideae) | Urophylleae | Antonio 1697 (AAU) | | Panama | de novo |
| BA031 | Urophyllum arboreum (Reinw. ex Blume) Korth. | Rubiaceae (Rubioideae) | Urophylleae | Boeea 7887 (S) | 1935 | Sumatra | RG-only |

*See the main text for details

**ENA accessions (beginning with ERR) refer to raw sequence reads. The respective assembled plastomes of these samples are available in the Dryad Digital Repository (https://doi.org/10.5061/dryad.mpg4f4r67)

**S1 Table. Taxa, voucher/source, ENA/GenBank, assembly, and sequencing information for sequences used in this study.**

Taxa with fully assembled plastomes are indicated in grey.

| Lab ID | Species | Family (Subfamily) | Tribe (Rubiaceae only) | Assembly reference seq | Plastome coverage | # Reads after dedupe and trimming | ENA/GenBank accesions** |
|---|---|---|---|---|---|---|---|
| | Alstonia scholaris (L.) R.Br. | Apocynaceae | | MN176280 | 12.0 | 3666186 | ERR5034019 |
| | Chironia baccifera L. | Gentianaceae | | ON641347 | 10.4 | 1484028 | ERR5033620 |
| | Geniostoma borbonicum (Lam.) Spreng. | Loganiaceae | | MT471262 | 72.2 | 5420570 | ERR5034022 |
| BD003 | Acranthera grandiflora Bedd. | Rubiaceae | Coptosapelteae | KY378704 | 41.4 | 9044646 | ERR9883540 |
| AY008 | Coptosapelta diffusa (Champ. ex Benth.) Steenis | Rubiaceae | Coptosapelteae | KY378704 | 180.9 | 8909362 | ERR9883544 |
| AZ040 | Luculia pinceana Hook. | Rubiaceae | Luculieae | NA | 134.5 | 11582072 | ERR9883503 |
| DE068 | Cinchona calisaya Wedd. | Rubiaceae (Cinchonoideae) | Cinchoneae | MZ151891 | 224.2 | 22815466 | ERR9883472 |
| CC071 | Deppea grandiflora Schltdl. | Rubiaceae (Cinchonoideae) | Hamelieae | KY378675 | 22.7 | 11600302 | ERR9883535 |
| DE066 | Paracorynanthe antankarana Capuron ex J.-F.Leroy | Rubiaceae (Cinchonoideae) | Hymenodictyeae | KY378679 | 63.8 | 29947254 | ERR9883470+ERR9883475 |
| | Corynanthe pachyceras K.Schum. | Rubiaceae (Cinchonoideae) | Naucleeae | KY378678 | 3.7 | 2323758 | ERR5033631 |
| AG098 | Rondeletia intermixta Britton | Rubiaceae (Cinchonoideae) | Rondeletieae | KY378681 | 80.8 | 19596792 | ERR9883534 |
| AP035 | Airosperma vanuense S.P.Darwin | Rubiaceae (Ixoroideae) | Airospermeae | KY348840 | 60.8 | 9756614 | ERR9883530 |
| | Razafimandimbisonia minor (Baill.) Kainul. & B.Bremer | Rubiaceae (Ixoroideae) | Alberteae | KY348839 | 7.9 | 1912346 | ERR5033626 |
| BE034 | Wendlandia dasythyrsa Miq. | Rubiaceae (Ixoroideae) | Augusteae | KY492076 | 100.5 | 42365064 | ERR9883548 |
| BD032 | Alseis lugonis L.Andersson | Rubiaceae (Ixoroideae) | Condamineeae | NC_036300 | 29.5 | 12403114 | ERR9883484 |
| | Emmenopterys henryi Oliv. | Rubiaceae (Ixoroideae) | Condamineeae | NC_036300 | 10.5 | 1894852 | ERR5033635 |
| CX014 | Ixora javanica (Blume) DC. | Rubiaceae (Ixoroideae) | Ixoreae | KY378663 | 67.1 | 24509876 | ERR9883554 |
| | Jackiopsis ornata (Wall.) Ridsdale | Rubiaceae (Ixoroideae) | Jackieae | KY378669 | 4.3 | 2271120 | ERR5033822 |
| | Heinsia crinita (Wennberg) G.Taylor | Rubiaceae (Ixoroideae) | Mussaendeae | KY348834 | 3.3 | 978442 | ERR5033638 |
| AV015 | Molopanthera paniculata Turcz. | Rubiaceae (Ixoroideae) | Posoquerieae | NA | 74.0 | 21299400 | ERR9883549 |
| CG080 | Sabicea marojejyensis Razafim. & J.S.Mill. | Rubiaceae (Ixoroideae) | Sabiceeae | NA | 357.2 | 18732102 | ERR9883531 |
| | Cyclophyllum barbatum (G.Forst.) N.Hallé & J.Florence | Rubiaceae (Ixoroideae) | Vanguerieae | KY378666 | 57.3 | 10233994 | ERR5034026 |
| CA098 | Glionnetia sericea (Baker) Tirveng. | Rubiaceae (Ixoroideae) | | NA | 111.4 | 8109216 | ERR9883500 |
| DE070 | Anthospermum thymoides Baker | Rubiaceae (Rubioideae) | Anthospermeae | CY051: this study | 26.8 | 12886590 | ERR9883532 |
| DE067 | Carpacoce spermacocea (Rchb. ex Spreng.) Sond. | Rubiaceae (Rubioideae) | Anthospermeae | CY051: this study | 913.3 | 23868312 | ERR9883471 |
| | Coprosma longifolia A.Gray | Rubiaceae (Rubioideae) | Anthospermeae | CY003: this study | 29.7 | 3417710 | ERR5034025 |
| CY012 | Coprosma rhamnoides A.Cunn. | Rubiaceae (Rubioideae) | Anthospermeae | CY003: this study | 144.9 | 13855942 | ERR9883542 |
| DE081 | Durringtonia paludosa R.J.F.Hend. & Guymer | Rubiaceae (Rubioideae) | Anthospermeae | CY003: this study | 36.0 | 9797814 | ERR9883564 |
| CX098 | Galopina aspera (Eckl. & Zeyh.) Walp. | Rubiaceae (Rubioideae) | Anthospermeae | CY051: this study | 46.6 | 8056560 | ERR9883565 |
| CX099 | Leptostigma pilosum (Benth.) Fosberg | Rubiaceae (Rubioideae) | Anthospermeae | CY003: this study | 200.0 | 19283612 | ERR9883489+ERR11475413 |
| CY003 | Nertera dichondrifolia (A.Cunn.) Hook.f. | Rubiaceae (Rubioideae) | Anthospermeae | NA | 108.9 | 9800008 | ERR9883499 |
| DE075 | Normandia neocaledonica Hook.f. | Rubiaceae (Rubioideae) | Anthospermeae | CY003: this study | 206.9 | 13472450 | ERR9883512 |
| CY089 | Opercularia volubilis R.Br. ex Benth. | Rubiaceae (Rubioideae) | Anthospermeae | NA | 268.1 | 11482826 | ERR9883537 |
| CY051 | Phyllis nobla L. | Rubiaceae (Rubioideae) | Anthospermeae | NA | 1051.4 | 14189504 | ERR9883493 |
| DE074 | Pomax umbellata (Gaertn.) Sol. ex A.Rich. | Rubiaceae (Rubioideae) | Anthospermeae | CY003: this study | 360.2 | 16524524 | ERR9883546 |
| BL040 | Argostemma elatostemma Hook.f. | Rubiaceae (Rubioideae) | Argostemmateae | CD007: this study | 73.8 | 12664432 | ERR9883520 |
| BU095 | Clarkella nana (Edgew.) Hook.f. | Rubiaceae (Rubioideae) | Argostemmateae | CD007: this study | 11.7 | 6379438 | ERR9883563 |
| CH091 | Mouretia larsenii Tange | Rubiaceae (Rubioideae) | Argostemmateae | NA | 171.9 | 18240426 | ERR9883550 |
| CD007 | Mycetia bracteata Hutch. | Rubiaceae (Rubioideae) | Argostemmateae | NA | 112.1 | 10711816 | ERR9883494 |
| | Mycetia Reinw. | Rubiaceae (Rubioideae) | Argostemmateae | CD007: this study | 15.9 | 1893208 | ERR5084289 |
| BK078 | Neohymenopogon parasiticus (Wall.) Bennet | Rubiaceae (Rubioideae) | Argostemmateae | CD007: this study | 100.1 | 7139796 | ERR9883558 |
| BD002 | Colletoecema dewevrei (De Wild.) E.M.A.Petit | Rubiaceae (Rubioideae) | Colletoecemateae | NA | | NA | KY378707 |
| BA061 | Coccocypselum condalia Pers. | Rubiaceae (Rubioideae) | Coussareeae | KY378701 + AY100: this study | 376.5 | 12339184 | ERR9883533 |

**S1 Table. Taxa, voucher/source, ENA/GenBank, assembly, and sequencing information for sequences used in this study.**

Taxa with fully assembled plastomes are indicated in grey.

| Lab ID | Species | Family (Subfamily) | Tribe (Rubiaceae only) | Assembly reference seq | Plastome coverage | # Reads after dedupe and trimming | ENA/GenBank accesions** |
|---|---|---|---|---|---|---|---|
| DA042 | Cruckshanksia pumila Clos in C.Gay | Rubiaceae (Rubioideae) | Coussareeae | KY378701 + AY100: this study | 2259.7 | 17990498 | ERR9883536 |
| DB007 | Faramea multiflora A.Rich. ex DC. | Rubiaceae (Rubioideae) | Coussareeae | KY378701 + AY100: this study | 167.6 | 27105744 | ERR9883469+ERR9883477 |
| | Craterispermum Benth. 2 | Rubiaceae (Rubioideae) | Craterispermeae | KY378698 | 14.5 | 2256818 | ERR5084275 |
| CK013 | Craterispermum Benth. 1 | Rubiaceae (Rubioideae) | Craterispermeae | KY378698 | 123.5 | 7060026 | ERR9883485 |
| CL038 | Craterispermum schweinfurthii Hiern | Rubiaceae (Rubioideae) | Craterispermeae | KY378698 | 95.3 | 7542388 | ERR9883498 |
| BZ084 | Cyanoneuron cyaneum (Hallier f.) Tange | Rubiaceae (Rubioideae) | Cyanoneuroneae | MN883829 | 89.2 | 12785706 | ERR9883481 |
| | Cyanoneuron pedunculatum Tange | Rubiaceae (Rubioideae) | Cyanoneuroneae | BZ084: this study | 0.5 | 676640 | ERR5034803 |
| DB070 | Danais nigra Homolle | Rubiaceae (Rubioideae) | Danaideae | DB077:this study | 171.0 | 18501058 | ERR9883478 |
| DB077 | Payera decaryi (Homolle) Buchner & Puff | Rubiaceae (Rubioideae) | Danaideae | NA | 148.3 | 26955428 | ERR9883539 |
| DC094 | Schismatoclada marojejyensis Humbert | Rubiaceae (Rubioideae) | Danaideae | NA | 479.4 | 22354314 | ERR9883567 |
| BC005 | Dunnia sinensis Tutcher | Rubiaceae (Rubioideae) | Dunnieae | MN883829 | 509.7 | 14233770 | ERR9883505 |
| CI009 | Gaertnera obovata Baker | Rubiaceae (Rubioideae) | Gaertnereae | KY378695 | 242.3 | 5111686 | ERR9883479 |
| | Gaertnera rotundifolia Bojer | Rubiaceae (Rubioideae) | Gaertnereae | KY378695 | 5.2 | 842634 | ERR5033654 |
| CA030 | Pagamea capitata Benth. | Rubiaceae (Rubioideae) | Gaertnereae | KY378695 | 24.7 | 10429052 | ERR9883502 |
| AP063 | Otiophora caerulea (Hiern) Bullock | Rubiaceae (Rubioideae) | Knoxieae | AI014: this study | 116.2 | 8571588 | ERR9883525 |
| AI014 | Chamaepentas hindsioides (K.Schum.) Kårehed & B.Bremer | Rubiaceae (Rubioideae) | Knoxieae | NA | 314.4 | 10619468 | ERR9883496 |
| DE064 | Triainolepis xerophila (Bremek.) Kårehed & B.Bremer | Rubiaceae (Rubioideae) | Knoxieae | AI014: this study | 983.5 | 50239610 | ERR9883541 |
| CQ058 | Lasianthus Jack | Rubiaceae (Rubioideae) | Lasiantheae | CH072: this study | 41.4 | 16361860 | ERR9883556 |
| CH072 | Lasianthus strigosus Wight | Rubiaceae (Rubioideae) | Lasiantheae | KY378708 | 86.0 | 10421850 | ERR9883504 |
| AZ001 | Ronabea latifolia Aubl. | Rubiaceae (Rubioideae) | Lasiantheae | CH072: this study | 308.4 | 9834184 | ERR9883501 |
| | Saldinia aegialodes Bremek. | Rubiaceae (Rubioideae) | Lasiantheae | AA052: this study | 10.6 | 1649566 | ERR5033656 |
| AA052 | Saldinia pallida Bremek. | Rubiaceae (Rubioideae) | Lasiantheae | AY100: this study | 3841.1 | 61673872 | ERR9883473+ERR9883476 |
| BE046 | Trichostachys microcarpa K.Schum. | Rubiaceae (Rubioideae) | Lasiantheae | AY100: this study | 109.1 | 11768826 | ERR9883545 |
| BZ092 | Mitchella repens L. | Rubiaceae (Rubioideae) | Mitchelleae | NA | | NA | KY378710 |
| AQ075 | Appunia guatemalensis Donn.Sm. | Rubiaceae (Rubioideae) | Morindeae | CV018: this study | 88.8 | 5879036 | ERR9883516 |
| | Coelospermum paniculatum F.Muell. | Rubiaceae (Rubioideae) | Morindeae | CV018: this study | 18.7 | 1270746 | ERR5034257 |
| BZ099 | Gynochthodes officinalis (F.C.How) Razafim. & B.Bremer | Rubiaceae (Rubioideae) | Morindeae | CV018: this study | 178.2 | 19554914 | ERR9883527 |
| CV018 | Morinda citrifolia L. | Rubiaceae (Rubioideae) | Morindeae | NA | 693.4 | 23023506 | ERR9883553 |
| AX046 | Lerchea bracteata Valeton | Rubiaceae (Rubioideae) | Ophiorrhizeae | MW528277 | 127.6 | 28155166 | ERR9883566 |
| | Kajewskiella trichantha Merr. & L.M.Perry | Rubiaceae (Rubioideae) | Ophiorrhizeae | MW528277 | 3.5 | 424886 | ERR5034845 |
| CH086 | Neurocalyx zeylanicus Hook. | Rubiaceae (Rubioideae) | Ophiorrhizeae | MW528277 | 147.7 | 16014084 | ERR9883555 |
| CY100 | Ophiorrhiza darwinii Razafim. & Rydin | Rubiaceae (Rubioideae) | Ophiorrhizeae | MW528277 | 69.9 | 13073088 | ERR9883497 |
| CZ012 | Ophiorrhiza mungos L. | Rubiaceae (Rubioideae) | Ophiorrhizeae | MW528277 | 177.9 | 10451454 | ERR9883526 |
| | Ophiorrhiza winkleri Valeton | Rubiaceae (Rubioideae) | Ophiorrhizeae | MW528277 | 5.1 | 852822 | ERR5033642 |
| | Paederia thouarsiana Baill. | Rubiaceae (Rubioideae) | Paederieae | NC_049155 | 34.4 | 2389358 | ERR5033643 |
| P0085 | Leptodermis potaninii Batalin | Rubiaceae (Rubioideae) | Paederieae | NC_049155 | 200.1 | 15133514 | ERR9883547 |
| CA022 | Paederia ciliata (Bartl. ex DC.) Standl. | Rubiaceae (Rubioideae) | Paederieae | NC_049155 | 92.0 | 12507838 | ERR9883513 |
| BX093 | Pseudopyxis heterophylla (Miq.) Maxim. | Rubiaceae (Rubioideae) | Paederieae | NC_049155 | 49.4 | 22257894 | ERR9883524 |
| C0005 | Serissa foetida (L.f.) Lam. | Rubiaceae (Rubioideae) | Paederieae | NC_049155 | 180.8 | 15697518 | ERR9883487 |
| B0110 | Spermadictyon suaveolens Roxb. | Rubiaceae (Rubioideae) | Paederieae | NC_049155 | 183.4 | 12712938 | ERR9883508 |
| AE034 | Rudgea recurva Müll.Arg. | Rubiaceae (Rubioideae) | Palicoureeae | KY378697 | 22.1 | 7703636 | ERR9883488 |
| BH070 | Palicourea alpina (Sw.) DC. | Rubiaceae (Rubioideae) | Palicoureeae | KY378697 | 42.4 | 17823444 | ERR9883557 |
| | Palicourea nitidella (Müll.Arg.) Standl. | Rubiaceae (Rubioideae) | Palicoureeae | KY378697 | 7.0 | 954302 | ERR5033644 |

**S1 Table. Taxa, voucher/source, ENA/GenBank, assembly, and sequencing information for sequences used in this study.**

Taxa with fully assembled plastomes are indicated in grey.

| Lab ID | Species | Family (Subfamily) | Tribe (Rubiaceae only) | Assembly reference seq | Plastome coverage | # Reads after dedupe and trimming | ENA/GenBank accesions** |
|---|---|---|---|---|---|---|---|
| CM050 | Puffia gerrardii (Baker) Razafim. & B.Bremer | Rubiaceae (Rubioideae) | Palicoureeae | KY378697 | 53.4 | 6652086 | ERR9883480 |
| | Perama dichotoma Poepp. | Rubiaceae (Rubioideae) | Perameae | AM028: this study | 9.9 | 1938690 | ERR5033646 |
| AM028 | Perama hirsuta Aubl. | Rubiaceae (Rubioideae) | Perameae | NA | 284.7 | 24301482 | ERR9883561 |
| CA031 | Prismatomeris fragrans E.T.Geddes | Rubiaceae (Rubioideae) | Prismatomerideae | NA | NA | NA | KY378699 |
| | Prismatomeris Thwaites | Rubiaceae (Rubioideae) | Prismatomerideae | KY378699 | 6.3 | 1217278 | ERR5084277 |
| CB078 | Rennellia subsessilis (King & Gamble) Razafim. & Rydin | Rubiaceae (Rubioideae) | Prismatomerideae | KY378699 | 367.2 | 16352526 | ERR9883521 |
| AF076 | Psychotria ankarensis (Bremek.) Razafim. & B.Bremer | Rubiaceae (Rubioideae) | Psychotrieae | NA | 99.6 | 3212850 | ERR9883486 |
| AG024 | Psychotria mahonii C.H.Wright | Rubiaceae (Rubioideae) | Psychotrieae | KY378696 | 33.4 | 5238420 | ERR9883483 |
| CL028 | Calycosia lageniformis (Gillespie) A.C.Sm. | Rubiaceae (Rubioideae) | Psychotrieae | AF076:this study | 103.6 | 5095646 | ERR9883559 |
| | Calycosia petiolata A.Gray | Rubiaceae (Rubioideae) | Psychotrieae | CL028: this study | 5.6 | 1412462 | ERR5034836 |
| | Chaetostachydium barbatum Ridsdale | Rubiaceae (Rubioideae) | Psychotrieae | CL028: this study | 2.0 | 670082 | ERR5034846 |
| | Dolianthus montiswilhelmii (P.Royen) A.P.Davis | Rubiaceae (Rubioideae) | Psychotrieae | AF076:this study | 28.9 | 3568582 | ERR5034835 |
| | Psychotria pandurata Verdc. | Rubiaceae (Rubioideae) | Psychotrieae | KY378696 | 1.5 | 778050 | ERR5033649 |
| | Plocama calabrica (L.f.) M.Backlund & Thulin | Rubiaceae (Rubioideae) | Putorieae | KY378690 | 8.2 | 1461868 | ERR5033647 |
| AH008 | Plocama tinctoria (Balf.f.) M.Backlund & Thulin | Rubiaceae (Rubioideae) | Putorieae | KY378690 | 346.4 | 18586312 | ERR9883529 |
| AH009 | Plocama dubia (Aitch. & Hemsl.) N. Backlund & Thulin | Rubiaceae (Rubioideae) | Putorieae | KY378690 | 58.4 | 6150778 | ERR9883507 |
| | Rubia peregrina L. | Rubiaceae (Rubioideae) | Rubieae | NC_047470 | 4.2 | 3603986 | ERR5033650 |
| AG064 | Sherardia arvensis L. | Rubiaceae (Rubioideae) | Rubieae | NC_047470 | 389.1 | 21195568 | ERR9883522 |
| M0004 | Didymaea alsinoides (Schltdl. & Cham.) Standl. | Rubiaceae (Rubioideae) | Rubieae | NC_047470 | 92.0 | 10419852 | ERR9883528 |
| DE065 | Galium polyacanthum (Baker) Puff | Rubiaceae (Rubioideae) | Rubieae | NC_047470 | 84.6 | 6173124 | ERR9883538 |
| T0054 | Kelloggia galioides Torr. | Rubiaceae (Rubioideae) | Rubieae | NA | 2114.7 | 38841036 | ERR9883509+ERR9883560 |
| BO005 | Rubia cordifolia subsp. conotricha (Gand.) Verdc. | Rubiaceae (Rubioideae) | Rubieae | NC_047470 | 302.4 | 32273554 | ERR9883467+ERR9883474 |
| CH081 | Schizocolea linderi (Hutch. & Dalziel) Bremek. | Rubiaceae (Rubioideae) | Schizocoleeae | NA | | NA | KY378700 |
| BZ091 | Lecananthus erubescens Jack | Rubiaceae (Rubioideae) | Schradereae | CA038: this study | 55.8 | 11232010 | ERR9883543 |
| CX040 | Schradera nervulosa (Stapf) Puff, R.Buchner & Greimler | Rubiaceae (Rubioideae) | Schradereae | CA038: this study | 212.2 | 20908840 | ERR9883523 |
| CA038 | Schradera rotundata Standl. ex Steyerm. | Rubiaceae (Rubioideae) | Schradereae | NA | 329.0 | 8213208 | ERR9883510 |
| CM080 | Seychellea sechellarum (Baker) Razafim., Kainul. & Rydin | Rubiaceae (Rubioideae) | Seychelleeae | KY378707 | 74.3 | 7073510 | ERR9883491 |
| CG025 | Diodella sarmentosa (Sw.) Bacigalupo & E. L. Cabral | Rubiaceae (Rubioideae) | Spermacoceae | DE079:this study | 81.1 | 7522408 | ERR9883492 |
| BZ040 | Exallage chrysotricha (Palib.) Neupane & N.Wikstr. | Rubiaceae (Rubioideae) | Spermacoceae | DE079:this study | 84.7 | 8374914 | ERR9883482 |
| DE079 | Oldenlandia herbacea (L.) Roxb. | Rubiaceae (Rubioideae) | Spermacoceae | NA | 1079.9 | 28408956 | ERR9883552 |
| | Spermacoce L. | Rubiaceae (Rubioideae) | Spermacoceae | DE079:this study | 9.6 | 1180896 | ERR5084278 |
| | Theligonum cynocrambe L. | Rubiaceae (Rubioideae) | Theligoneae | KY378688 | 78.1 | 5963764 | ERR5033810 |
| BV076 | Theligonum japonicum Ôkubo & Makino | Rubiaceae (Rubioideae) | Theligoneae | KY378688 | 303.0 | 57226818 | ERR9883515+ERR9883518 |
| AX027 | Amphidasya longicalycina (Dwyer) C.M.Taylor | Rubiaceae (Rubioideae) | Urophylleae | AY100: this study | 63.2 | 7053006 | ERR9883495 |
| AS084 | Temnopteryx sericea Hook.f. | Rubiaceae (Rubioideae) | Urophylleae | AY100: this study | 70.8 | 9626114 | ERR9883562 |
| | Urophyllum cyphandrum Stapf | Rubiaceae (Rubioideae) | Urophylleae | AY100: this study | 4.0 | 1574358 | ERR5033613 |
| AY100 | Raritebe palicoureoides Wernham | Rubiaceae (Rubioideae) | Urophylleae | NA | 269.7 | 9986836 | ERR9883517 |
| BA031 | Urophyllum arboreum (Reinw. ex Blume) Korth. | Rubiaceae (Rubioideae) | Urophylleae | AY100: this study | 11.3 | 3468372 | ERR9883490 |

*See the main text for details

**ENA accessions (beginning with ERR) refer to raw sequence reads. The respective assembled plastomes of these samples are available in the Dryad Digital Repository (https://doi.org/10.5061/dryad.mpg4f4r67)

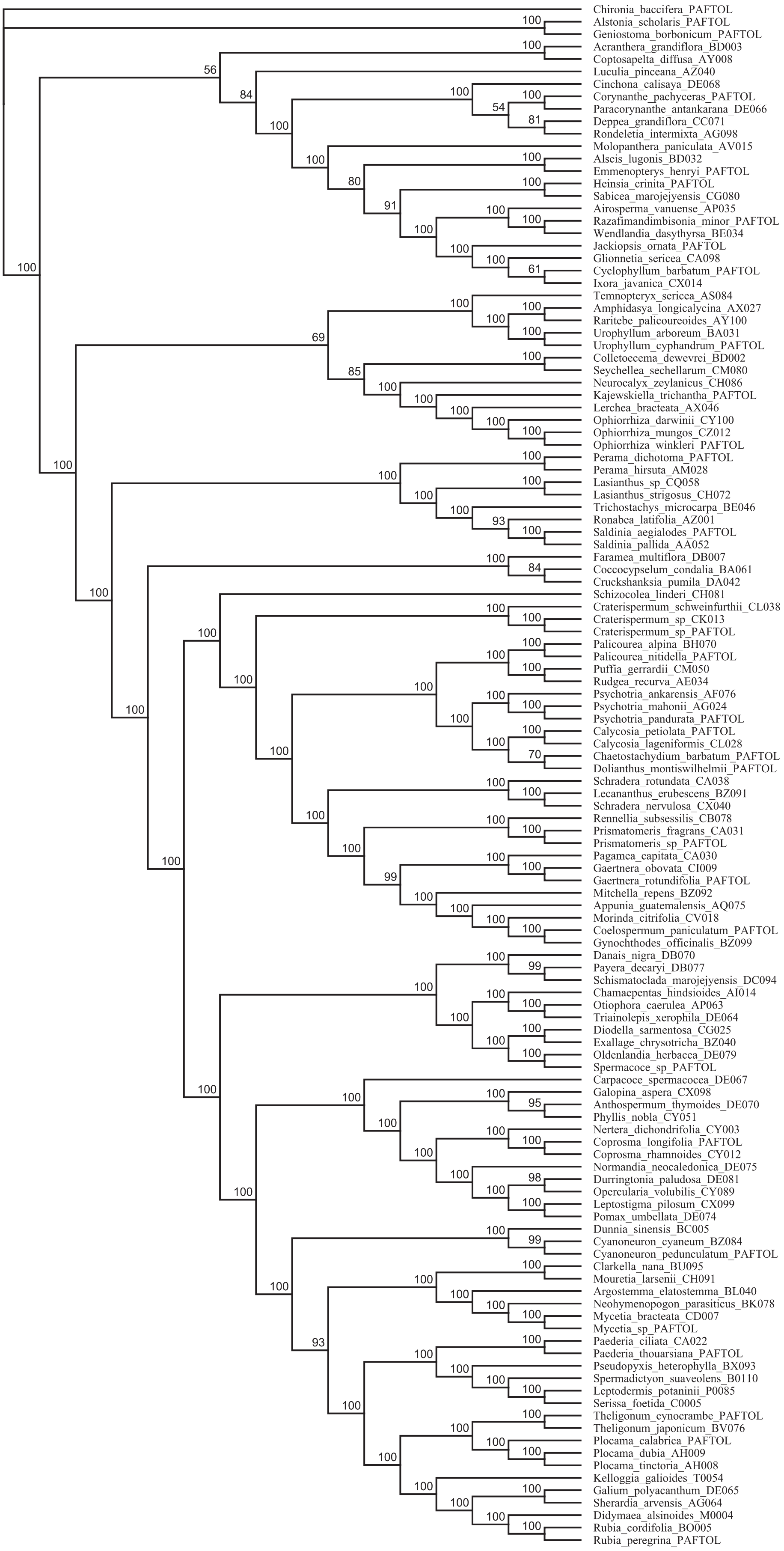


**S1 Fig. Plastome phylogeny inferred from maximum likelihood analysis of the RY-coded version of alignment 1.** The four nucleotides were recoded into two states (purines and pyrimidines). Alignment 1 was untrimmed except for removal of autapomorphies (columns where more than 99% of the samples had a gap). Numbers above branches indicate bootstrap support values.

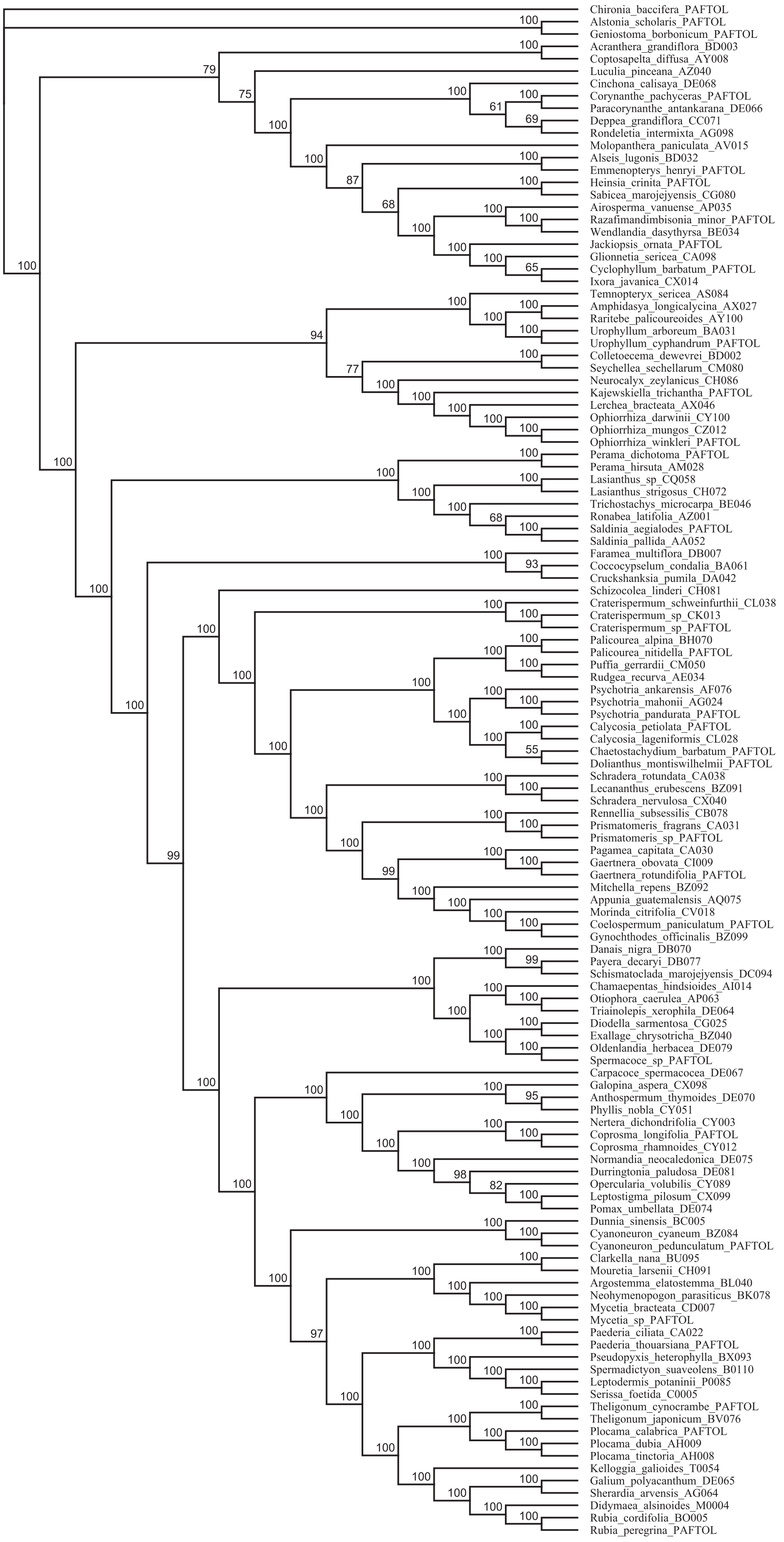


**S2 Fig. Plastome phylogeny inferred from maximum likelihood analysis of the RY-coded version of alignment 2.** The four nucleotides were recoded into two states (purines and pyrimidines). Alignment 2 was trimmed using trimAl. Numbers above branches indicate bootstrap support values.
